# Plant diversity promotes aboveground arthropods and associated functions despite arthropod loss over time

**DOI:** 10.1101/2025.04.09.647912

**Authors:** A. Ebeling, Maximilian Bröcher, Lionel Hertzog, Holger Schielzeth, Wolfgang W. Weisser, Sebastian T. Meyer

## Abstract

Arthropods comprise the majority of terrestrial biodiversity and play key roles in ecosystem functioning. Biodiverse grasslands support many arthropods, yet such ecosystems have declined due to land conversion and management changes. While restoration aims to conserve species-rich grasslands, long-term effects of plant richness on arthropod communities and associated functions remain underexplored. We addressed this gap by quantifying arthropods, herbivory, and predation over 13 years (2010-2022) across 80 grassland plots with varying plant richness. We examined (1) temporal trends in arthropod communities, herbivory and predation and whether changes depended on plant richness, (2) whether plant richness effects varied or strengthened over time, and (3) whether arthropod changes affected associated functions. Arthropod metrics declined over time across all plant richness levels, with average losses mainly being more pronounced in species-poor mixtures. Plant richness consistently had a positive effect on arthropods and their functions, although this effect varied between years without a consistent temporal trend. Notably, temporal changes in arthropod community metrics did not predict shifts in associated functions. Our findings highlight the dynamic interplay between plant richness and arthropods. From a conservation perspective, we can conclude that diversification in grasslands-the increase in plant diversity-directly supports arthropods and associated functions. Additionally, first trends indicate that the maintenance and protection of diverse semi-natural grasslands over a long period might mitigate the arthropod loss driven by environmental changes. In other words, diverse grasslands may buffer against the ongoing arthropod loss, though this effect may take years to become apparent. This again emphasizes the long-term nature of conservation efforts.

## Introduction

Arthropods make up about 96% of known biodiversity in terrestrial ecosystems. As *“the little things that run the world”* ^1^, they strongly contribute to the high social, cultural and economic value of many habitats ^2,3^. Arthropods also provide a variety of important ecosystem functions ^4–7^, such as decomposing dead plant material and enriching the soil with plant-available nutrients ^5,8^. As herbivores, they mediate the transport of energy from primary producers to higher trophic levels ^9^ and cause feedbacks to plant fitness and community composition ^10^. In addition to their role as herbivores, many arthropods also serve as pollinators, facilitating the reproduction and survival of various plant species ^11,12^, or as predators, regulating populations of herbivorous insects ^8,13^. Furthermore, arthropods build a fundamental food resource for other consumers, such as birds and many other vertebrates, thereby acting as an integral component of food webs ^14^. Given the crucial role of arthropods in many ecosystems, reports of drastic declines in their abundance, biomass and diversity have sparked both scientific and societal concern ^15,16^. Over the past decades, global arthropod declines have been well-documented, with biomass losses reaching up to 75% and significant reductions in species richness observed across diverse habitats, including grasslands^17–20^.

Grasslands cover about 40% of the earth’s terrestrial surface ^21^ and around 13% of Europe ^22^ (32 European countries included in this study) and are thus one of the most common ecosystems with important contributions to global ecosystem functioning and services ^3^. There are two main types of grasslands: natural grasslands, which remain intact without human intervention, and semi-natural grasslands, which require a minimum level of anthropogenic management to maintain an open habitat and prevent succession ^23,24^. Extensively managed, semi-natural grasslands, which are maintained by low levels of grazing or mowing (extensively managed), are among the most species-rich ecosystems in Europe ^25^. Over the past centuries extensively managed grasslands have declined significantly, primarily as a result of land conversion into arable land or land use intensification characterized by more fertilization and more grazing or more frequent mowing ^21,23,25,26^. To counteract this trend and to meet targets of the Convention of Biological Diversity (CBD), conservation efforts should focus on the maintenance of existing species-rich semi-natural grasslands (conservation) and the restoration of currently species-poor grasslands (diversification) ^2,4,27^.

There is increasing evidence that species-rich grasslands not only function better than low-diversity grasslands, with respect to a large number of ecosystem functions, but also, that the benefits gained by increasing plant species richness become stronger over time ^28^. This applies in particular to increased plant species richness effects on plant productivity, but also to other ecosystem functions ^6,29–31^. Hence, maintaining existing species-rich semi-natural grasslands as a conservation strategy is expected to have stronger positive effects than newly establishing such grasslands. So far, however, there are only few studies investigating temporal changes in the relationship between plant species richness and arthropod communities with their mediated functions ^6,32^. In other words, our understanding of temporal dynamics in grasslands lacks a multitrophic perspective ^28^.

Arthropods generally show a higher species richness, diversity and abundance in grasslands with higher plant species richness and are also more diverse in their interactions and their functional traits compared to low-diversity grasslands ^7,33–37^. However, plant species richness effects may vary or even become stronger over time either because arthropods perform better in species-rich plant communities, perform worse in species-poor plant communities, or a combination of both ^6^. Alternatively, higher plant species richness might mitigate the ubiquitous declines in arthropod numbers, making the losses less severe. Hence while, arthropod communities in species-poor and in species-rich grasslands both tend to depauperate over time, the performance losses are expected to be less pronounced in species-rich grasslands. So far, no studies have empirically tested this pattern in grasslands.

Multiple mechanisms can cause variation in the relationship between plant species richness and arthropod communities, such as changes in the plant community composition ^29,38^ or abiotic conditions. Specifically, due to community assembly processes and selection for specific plant phenotypes, the functional and genetic diversity and thus the complementarity among plants might increase over time in diverse plant communities ^29,31,39,40^. In addition, species-rich plant communities fluctuate less in productivity, structure, or soil temperature ^41–43^. Because arthropods in grasslands are strongly bottom-up controlled ^4,44,45^, plant species richness-induced temporal changes in plant attributes are expected to escalate to arthropod communities, their interactions and related processes ^28,36,46–48^. A greater arthropod diversity (as found in species-rich plant communities) can lead to greater temporal stability in arthropod communities and contribute to changes in the relationship between plant species richness and arthropods over time via various mechanisms.

A high diversity of arthropods might increase the chance of communities to include species that perform well under changing or harsh environmental conditions (such as dry summers). Furthermore, diverse arthropod communities are more likely to include species that are either very robust (trait-dependent resistance), have a high reproduction rate (fast recovery), or have capacity to adapt (highly plastic in their traits). These effects are known as sampling or selection effect. Alternatively, or in addition, a high functional complementarity between arthropod species combined with asynchronous fluctuations (asynchrony) could lower fluctuations in arthropod community properties and arthropod-mediated functions (portfolio or insurance effect of biodiversity) ^49–51^.

Despite these insights, the enormous diversity, and the critical ecosystem functions provided by arthropods in grasslands, there has been no long-term study on arthropod communities in plant diversity experiments exploring the temporal trajectory of cascading biodiversity effects. To address this gap, we here test how experimentally manipulated plant species richness influences arthropod communities, plant consumption by herbivores, and predation in a semi-natural grassland over a time course of 13 years. Using standardized methods, we collected arthropods, measured leaf damage and quantified predation on 80 experimental plots differing in plant species richness. We used these data to calculate complementary arthropod community metrics (species richness, diversity, abundance, biomass for all arthropods together and herbivores and predators separately) and their mediated functions (biomass ratios across trophic groups, percentage herbivory and predation rate). Whereas arthropod community metrics inform about the “attractiveness” of plant communities for arthropods, arthropod related ecosystem functions allow us to quantify top-down effects on plant communities and herbivores ^13,35^. With this approach, we addressed the following questions:

1. Are there temporal trends in arthropod communities and associated functions (declines or gains over the study period), and are these changes influenced by plant species richness? Based on the reported global loss in arthropods and the expected buffering effect of plant species richness, we predict that overall arthropod communities will decline, but that losses will be less pronounced in species-rich plant communities than in species-poor ones.
2. Do effects of plant species richness on arthropod communities vary or even strengthen over time? Based on the finding that the relationship between plant species richness and productivity becomes stronger over time and the potentially buffering effect of species richness, we expect a strengthening plant species richness effect over time.
3. Do temporal changes in the strength of the relationship between plant species richness and arthropod communities affect associated functions, such as the biomass ratios between arthropods and plants, herbivores and plants, predators and herbivores, herbivory and predation? We expect a strong coupling of stocks and flows, so that the performance of arthropod communities explains the temporal variation in predation and herbivory.

To answer those questions, we applied three consecutive models, which separately tested for temporal trends and its dependence on plant species richness (model 1), variation in the effects of plant species richness (model 2) and strengthening of plant diversity effects over time (model 3).

## Results

In total, we collected about 44.900 individual arthropods assigned to 400 taxa. Coleoptera were the most dominant group, contributing 35% and 32% to overall abundance and species richness, respectively, followed by Hemiptera (33%, 34%), Hymenoptera (24%, 20%) and Araneae (8%, 14%). Overall, the collected arthropods were predominantly herbivores (66.7%) and predators (31.8%).

### Temporal trends in arthropod community metrics and associated functions and its dependence on plant species richness

Over a period of 11 years, we observed a strong decrease in all arthropod community metrics (Figures 1, S1, Tables S1, S2). Across seasons, at intermediate plant species richness, arthropod species richness and biomass decreased by −5.2% and −7.4% per year, respectively, with stronger decreases in spring than in summer (Table S2). When comparing herbivorous and predatory arthropods, we found that the annual losses were more pronounced in predators (Figure 1, S1, Table S2; richness: −6.1% versus −3.9%; biomass: −8.1% versus −7.0%). However, the annual rate of change differed slightly between high and low diverse plant communities and between sampling seasons (Figure 2, Table S2). For example, across sampling seasons, we found that the annual abundance loss of herbivores and predators was stronger in species-rich compared to species-poor plant communities (herbivore abundance high and low plant species richness: −6.5% versus −6.0%, predator abundance high and low plant species richness: −7.5% versus −6.3%), while we observed the opposite pattern for the annual loss in arthropod species richness (herbivore richness high and low plant species richness: −3.7% and −4.2%, predator richness high and low plant species richness: −6.0% versus −6.2%) (Figure 2, Table S2). Furthermore, in spring (May) the annual decrease in arthropod species richness, abundance and biomass was generally stronger in high species mixtures (60 plant species) compared to monocultures (except for predator richness). In contrast, during summer (July), herbivore (but not predator) loss in monocultures was more pronounced than the loss in high diverse plant communities (Figure 2, Table S2).

**Figure 1:**
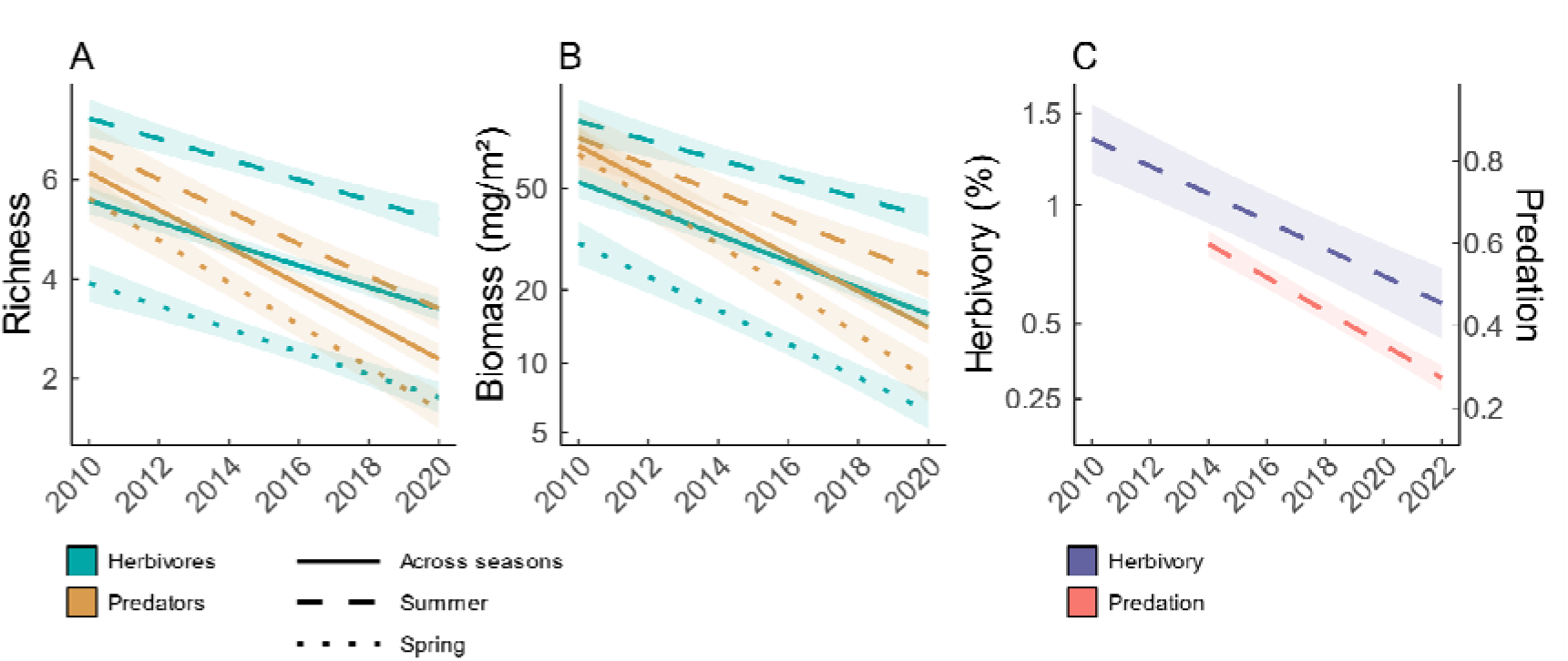
Temporal trends in arthropod richness, biomass and functions (herbivory and predation) Regression lines and 95% confidence intervals show the predicted relationship between time (in years) and arthropod richness (A), arthropod biomass (B), and arthropod related functions (C). Both were derived from model (1), where time was included as a numerical variable. Arthropods were sampled from 2010 to 2020, and are separated into herbivores (blue) and predators (orange). Herbivory (violet) and predation (red) were assessed from 2010 to 2022 and 2014 to 2022, respectively. Line styles indicate changes in spring (dotted), in summer (dashed) and across seasons (solid). Biomass and herbivory were log-transformed prior to the analyses.

**Figure 2:**
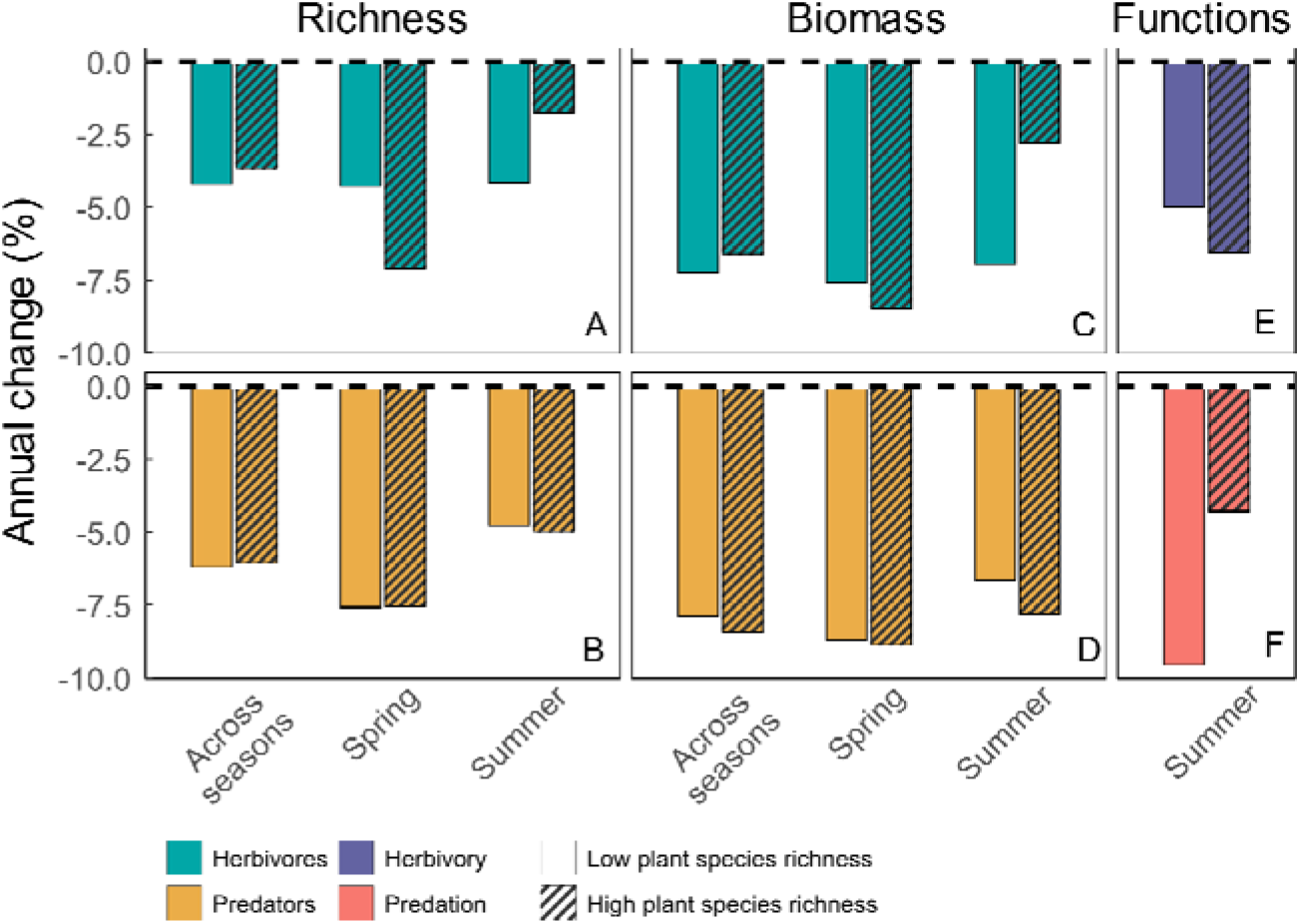
Annual (%) change in arthropod richness, biomass and functions (herbivory and predation). Bars show predicted annual changes in arthropod richness (A-B), arthropod biomass (C-D), herbivory (E) and predation (F). Values were extracted from model (1) for 60-species plant mixtures (plane bars) and monocultures (shaded bars), where time was included as a numerical variable (see Figure 1). Annual changes are shown separately for herbivores (blue), predators (orange), herbivory (violet) and predation (red), in spring, summer and across both seasons.

With respect to ecosystem functions, we observed a significant annual decline in herbivory (−4.8%) and predation (−9.1%), with comparably minor changes in the arthropod-plant ratio (−0.7%), and the predator-herbivore ratio (+0.4%) (Figures 1, 2, S2, Tables S3, S4). In contrast, the herbivore-plant ratio increased across season at intermediate level of plant species richness by +4.6% per year (Figure S2, Table S3). The annual rate of change varied between high-species richness plant communities (60 species) and monocultures. Predation loss was more pronounced in species-poor plant communities compared to species-rich ones (−11.9% versus −5.4%), whereas the opposite trend was observed for herbivory (−4.2% versus −5.5%) and the predator-herbivore biomass ratio (−0.7% versus −2.2%) (Figure 2, Table S3). Interestingly, the biomass ratios between arthropods and plants and herbivores and plants in high and low diverse plant communities showed completely opposing trends over time. Both biomass ratios showed strong annual increases in species-poor (arthropod-plant ratio +6.5%, herbivore-plant ratio +11.9%), but annual decreases in species rich plant communities (arthropod-plant ratio −7.8%, herbivore-plant ratio −7.3%) (Table S3).

### Effects of plant species richness on arthropod community metrics over time

Across all sampling years, we found a consistent positive effect of plant species richness on arthropod species richness, Hill number, abundance, and biomass—for the entire arthropod community, and for herbivores and predators separately (Figures 3, S3-S6, Table S5). However, for most of the metrics tested the effects of plant species richness varied significantly among years (Figures 3 S3-S6, Table S5, significant interactions between PSR and CY, and PSR, CY and Season). For example, the PSR slope ranged from 0.01 to 0.21 (PSR effects in 2020 and 2017, respectively) for predator richness in spring, and from 0.07 (2014) to 0.26 (PSR effects in 2019 and 2020, respectively) for predator richness in summer (Figure 4).

**Figure 3:**
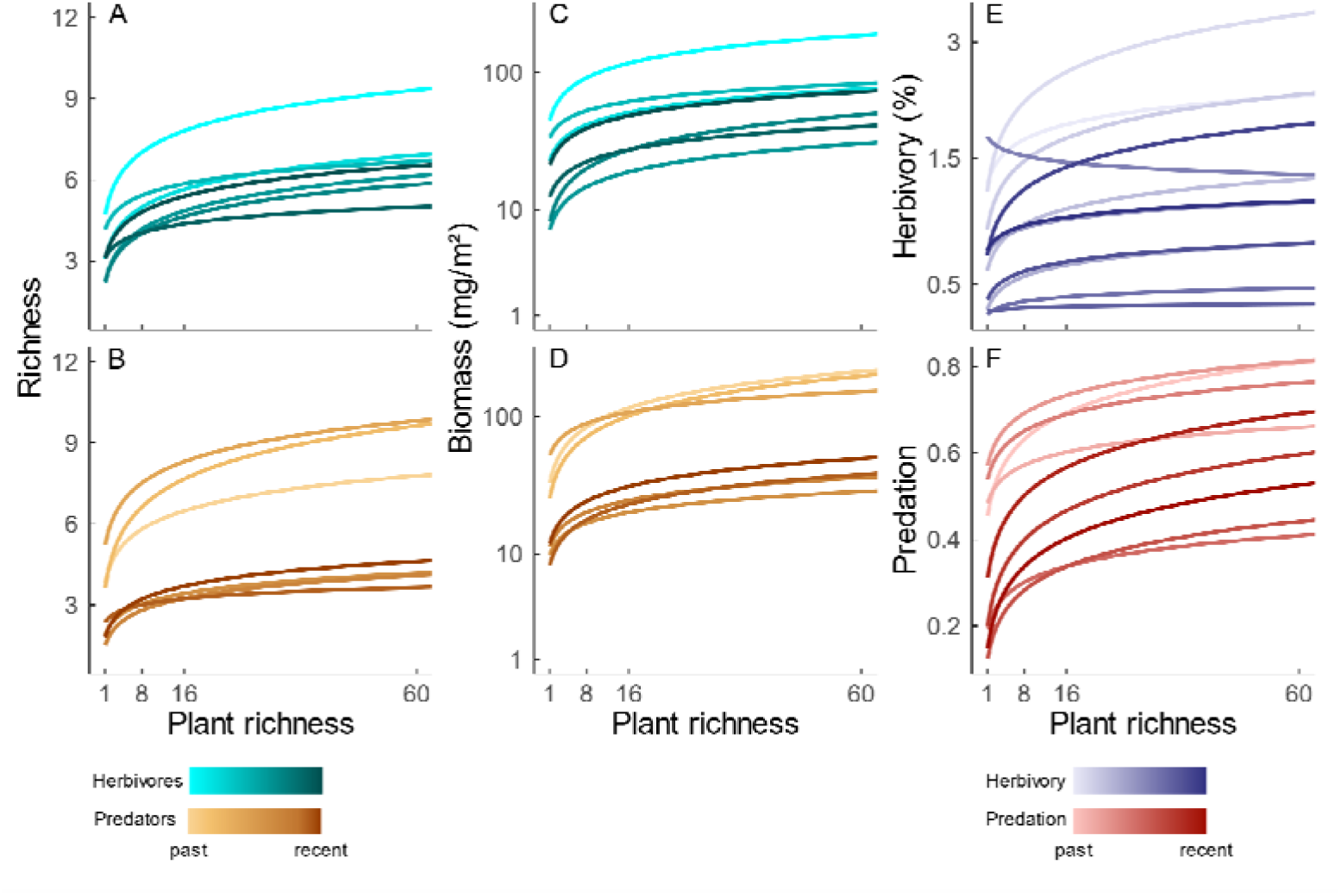
Plant species richness effects on arthropod richness, biomass and functions (herbivory and predation) in multiple years. Regression lines show the predicted relationship between plant species richness and arthropod richness (A-B), arthropod biomass (C-D), herbivory (E) and predation (F) for each sampling year. Regression lines were derived from model (2), where time was included as a factor, to allow for year-to year variation in the effects of plant species richness. Herbivores (blue) and predators (orange) were sampled between 2010 and 2020, and herbivory (violet) and predation (red) were assessed between 2010 and 2020, and 2014 and 2022, respectively. Biomass (mg/m²) and plant species richness were log-transformed prior to the analyses.

**Figure 4:**
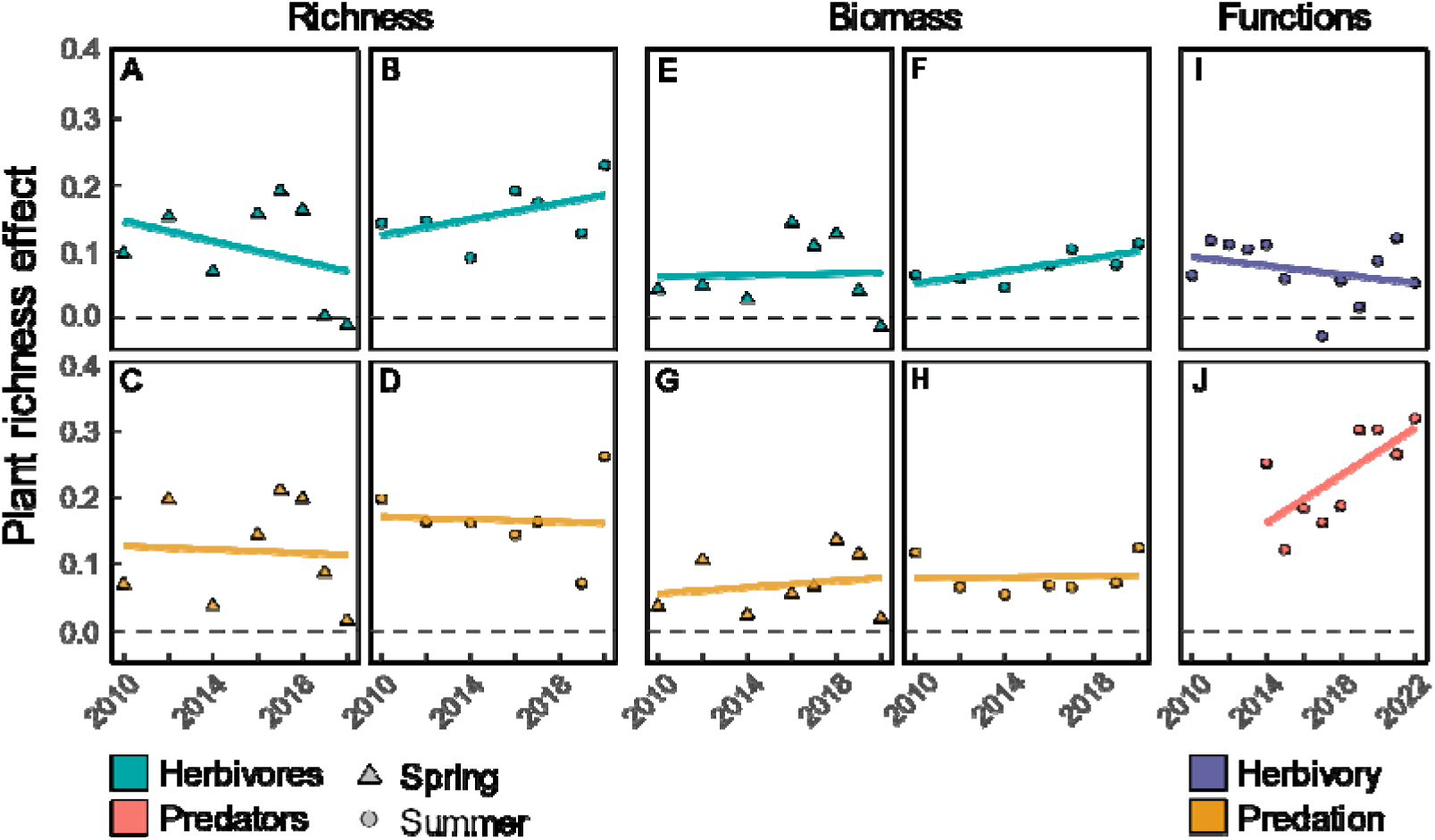
Change in the effect of plant species richness on arthropod richness, biomass and functions (herbivory and predation) over time. The regression lines and 95% confidence intervals show the predicted relationship between time (in years) and the effects of plant species richness on arthropod richness (A-D), arthropod biomass (E-G), and arthropod related functions (I-J). These were both derived from model (3), in which time was included as a numerical variable. To make the effects of plant species richness derived from model (2) comparable across years and independent of the absolute differences in arthropods and related functions between years (see Figure 3), the PSR slopes were corrected by dividing them by their respective yeaŕs average value. The data are shown for herbivores (blue) and predators (orange) in both seasons (triangles for spring and dots for summer).

When testing for directional temporal changes we found that the positive plant species richness effects on overall arthropod communities, herbivores and predators did not significantly change over time (Figures 4, S7, Tables 1, S6).

Arthropod communities and their responsiveness to changing plant species richness were also influenced by the sampling season. We consistently observed a significant increase in arthropod community metrics from spring to summer, with herbivore richness and biomass, for example, being around 134% and 178% higher in summer than in spring, respectively (Figures S4-S6, Table S5). Moreover, plant species richness effects on herbivore communities in summer tended to strengthen over time, whereas they either remained stable or weakened in spring communities (Figure 4, S7). Even though this is only significant for estimated richness and Hill number of herbivores (interaction of time and season in Tables 1 and S6), it is a consistent pattern for all metrics describing herbivore communities (Figures 4, S7).

### Effects of plant species richness on arthropod-mediated functions over time

Across all sampling years, we found a negative relationship between plant species richness and the overall arthropod-plant biomass ratio, as well as the biomass ratio between herbivores and plants (Figures S8, S9, Table S7). In other words, the amount of arthropod biomass (mg) per unit plant biomass (g) was generally lower in species-rich compared to species-poor plant communities. Contrary to what we might expect based on these findings, functions mediated by herbivores and predators (herbivory and predation) generally increased with increasing plant species richness, respectively (Figures 3 S10, Table S7).

The strength of the relationship between plant species richness and arthropod function varied significantly between years. For instance, the average plant species richness slopes for the arthropod-plant ratio in spring ranged from −0.08 (2012) to −0.40 (2020) (Figure S11) and the effect of plant species richness on herbivory ranged between slightly negative in a single year (−0.03 in 2017) to positive in all other years (maximum 0.12 in 2021) (Figures 4). Over time, the negative relationship between plant species richness and the biomass ratio of arthropods and plants and herbivores and plants became more pronounced, while the ratio between predator and herbivore biomass remained stable (Figures S11, Table S6). Furthermore, we observed an increase in the positive effect of plant species richness on predation, but not on herbivory (Figures 4, Table 1). In other words, the effect of plant species richness on predation strengthened over the 9-year period.

**Table 1:**
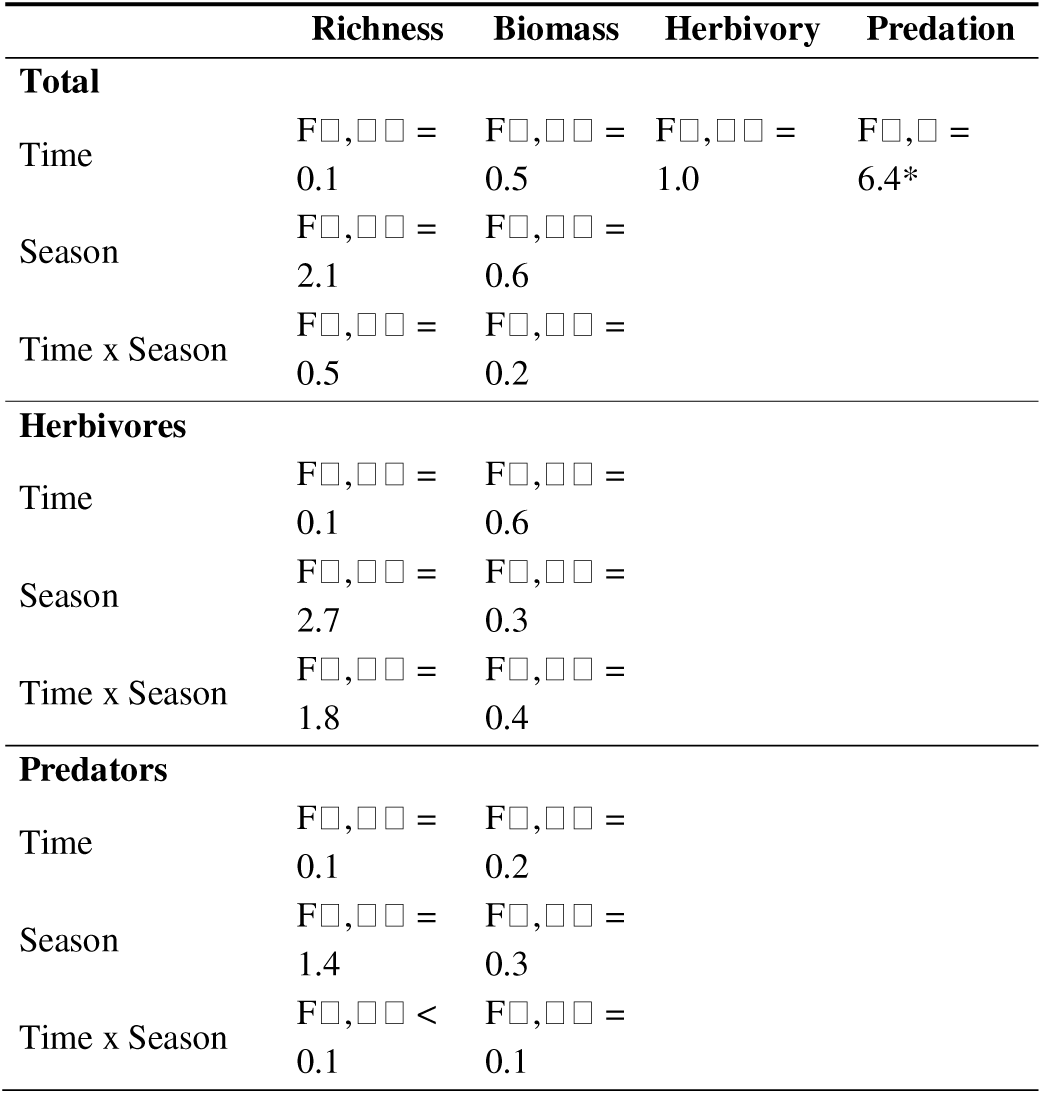
Changes in plant species richness effects over time. Results from linear models testing whether plant species richness effects on arthropod richness, biomass (mg/m²), herbivory and predation (PSR slope) change over **time** (calendar year as numerical variable), between sampling **seasons** (May/ spring and July/ summer) or for an interactive effect of both (**time x season**). For a detailed model description see model (3) in the section *Statistical analysis*. Significances are indicated by asterisks as followed: *** *P* < 0.001; ** *P* < 0.01; * *P* < 0.05.

## Discussion

Previous studies have shown strong decreases in arthropods during the last decades ^18,19,52,53^ and supporting effects of plant species richness for arthropods and associated functions ^4,33,35,54^. However, it is largely unknown whether plant species richness helps buffer arthropod loss over time and, as a consequence, how plant species richness effects might change over time ^28^. Based on our analysis of long-term arthropod, herbivory and predation data, we can draw conclusions about the interactive effect of plant species richness and time: (1) arthropod community properties strongly decrease over time, with declines being stronger in spring than in summer, (2) predators experience a higher annual loss than herbivores, (3) herbivore, but not predator loss is lower in species-rich plant mixtures in summer, but higher in spring, (4) the effects of plant species richness on arthropods are consistently positive, vary in strength among years, but neither consistently strengthen nor weaken over time, (5) plant species richness effects on predation strengthened over time, indicating that (6) temporal changes in arthropod community properties (stocks) in differently diverse plant communities do not predict the observed variation in arthropod associated functions (flows).

### Temporal trends in arthropod communities and its dependence on plant species richness

Over a period of 11 years, we could show a strong decline in arthropods, which aligns with previous studies documenting significant declines in arthropod richness, biomass and population size ^17–20^. With 76% loss between 2010 and 2020, the average annual loss in arthropod biomass in our study is higher than reported in many other studies. For example, in a study from 2017, Hallmann et al. ^17^ showed the similar total decline over a time period of 27 years, and in a meta-analysis, van Klink et al. ^19,20^ showed a decline in insect abundance of 11% in a decade. Most previous studies on arthropod decline cover data spanning several decades, but since the dramatic declines have only begun more recently, calculations based on longer time frames may show lower rates of average loss compared to those focused on the past few years ^19,20^. In addition, temporal dynamics in the plant or soil community that are independent of the reported global loss may also have caused the local decline in arthropods. Climatic changes or successional dynamics, for example, may have led to an overall decline in plant biomass, but also to changes in nutrient availability or soil conditions ^31,55^. Further mechanistic research is required to understand the role of successional dynamics in local arthropod loss. Annual loss in predatory arthropods was more pronounced than in herbivores, which can be explained by a higher sensitivity of higher trophic levels to abiotic changes ^56^. Interestingly, especially in herbivores and in summer communities, we observed slightly lower average annual losses in species-rich compared to species-poor plant mixtures, suggesting that the plant species richness effects could be strengthened over time ^6,28^.

### Inter-annual variation in the plant species richness-arthropod relationship

Our results consolidate the well-established link between high local plant species richness and a more complex arthropod community. Unlike previous studies, which frequently focused on particular aspects of the arthropod community and were limited to specific time points ^7,33,34,34,36,57,58^, our research demonstrates that this positive relationship applies to various facets of the arthropod community, remains consistent across seasons (spring and summer), and is stable over many years.

Mechanisms driving positive effects of plant species richness on arthropods include increased plant productivity, and structural complexity ^30^, which provide more and more diverse resources and microhabitats, supporting higher arthropod diversity and abundance ^59,60^. Since the described mechanisms are driven by bottom-up processes, we expected the positive relationship between plant species richness and arthropods to strengthen over time, similar to patterns observed for aboveground primary productivity ^29,31^. However, we found significant year-to-year variation in the strength of this relationship instead of a clear linear trend over time, with only a weak linear trend for individual arthropod community metrics (such as biomass or estimated richness). Overall, it appears that plant biomass alone does not drive the strength of the relationship between plant species richness and arthropod community metrics, and that other biotic (both extrinsic and intrinsic) and abiotic factors may play a crucial role. Potential candidates include interannual fluctuations in the effects of plant species richness on plant traits (such as defense traits and palatability) ^48,61,62^, plant species performance ^48^, or microclimatic conditions (such as soil moisture and temperature) ^42^. Furthermore, the stability of the arthropod community when faced with unfavorable conditions might also be an important factor. Various extreme events that occurred during the study period ^63^ have likely contributed to these fluctuations, making it impossible to see a clear linear trend over the 11-year period ^17^. Specifically, Müller et al. ^63^ showed that deviations from ‘optimal’ weather conditions significantly impact the population size of arthropods. Since the buffering effect of biodiversity (better performance of high-compared to low-species richness communities; ^38,49–51,64–66^) only comes into play under unfavorable conditions, climate extremes could contribute to the observed annual fluctuations in the plant species richness effect. In other words, to a certain extent, plant species richness effects on arthropod communities might be stronger under harsh conditions.

### Intra-annual variation among arthropod communities

Our results indicate that arthropod communities in spring and summer strongly differ in their response to time, plant species richness and their interactive effect. Arthropod communities appearing in spring experienced a higher annual loss than summer communities, maybe due to their weaker response to increasingly frequent winter weather anomalies ^63^. In contrast, due to their phenology, summer species may be better in coping with dry and hot conditions. Additionally, spring species might be more negatively affected by phenological shifts in plant communities or may experience a mismatch with their plant hosts due to earlier seasonal occurrences ^67,68^.

Our finding that loss in most arthropod community metrics is less pronounced in species-poor plant mixtures during spring and in species-rich plant mixtures during summer may be due to rising temperatures having a positive effect on arthropod communities in spring but a negative effect in summer. Specifically, as a consequence of spring temperatures rise, species-poor plant communities may heat up more than species-rich plant communities due to lower structural complexity. Consequently, since arthropod performance benefits from increasing temperature ^69^, the loss in monocultures could be lower than in high species mixtures. In summer, when temperatures are high, the more productive, species-rich mixtures are likely favored because they provide protection against heat, drought, and temperature fluctuations ^42^.

Finally, we could show that the positive relationship between plant species richness and arthropod community metrics was more pronounced in summer than in spring. Amongst others, this could be explained by the gradual development of plant species richness over the growing season; i.e. diverse plant mixtures become richer as the season progresses, leading to a cumulative effect on arthropod communities. Further, the better survival and accumulation of arthropods in species-rich plant mixtures might be more pronounced in summer, when species-poor plant mixtures become less favorable due to heat, resource limitations, and lack of shelter. In contrast, even in species poor plant mixtures, resources in spring are less limited, so the advantages of species-rich mixtures are less apparent early in the season.

Overall, plant species richness effects over time warrant further investigation, not only regarding the buffering effects of plant species richness across years but also within years among seasons. Future research should aim to identify the most critical time points during the vegetation period when arthropod losses are most pronounced.

### Links between temporal changes in arthropods (stocks) and associated functions (flows)

Our study reveals that temporal changes in arthropod community metrics differ from patterns observed for plant biomass in previous studies ^31^, both, in terms of the temporal trend and the dependency on plant species richness. Specifically, in plants, the decrease in biomass over time is stronger in monocultures than in species-rich plant communities ^31^, however, in arthropod communities the annual biomass loss across seasons is similar in high and low diverse mixtures (Figure 2C and D). As a consequence, the biomass ratio between arthropods and plants or herbivores and plants-increases over time in low and decreases in high plant species mixtures.

Further, responses in biomass to increasing plant species richness seem to differ in strength between arthropods and plants, i.e. plant biomass increases to a higher extent in species −rich compared to species-poor plant communities than arthropods, leading to a dilution effect in arthropod-plant and herbivore-plant ratios with increasing plant species richness ^35^. As the change in biomass ratios differs between high and low plant species richness, the negative effect of plant species richness becomes stronger over time (Figure S11).

Interestingly, as temporal changes and their response to changing plant species richness in herbivore and predator biomass follow similar patterns, the predator-herbivore ratio remains stable, both, over time and along the plant species richness gradient. This higher stability might indicate that predator biomass is even more bottom-up controlled (i.e. linked to its prey biomass) than herbivores are.

Surprisingly, neither temporal, nor plant species richness patterns for the directly assessed functions (herbivory and predation) can be explained by patterns in biomass ratios. Based on the biomass ratios, no loss and no plant species richness effect would be expected for predation. Similarly, for herbivory, patterns in biomass ratios suggest a negative relationship between plant species richness and herbivory and a decreasing plant species richness-herbivory relationship over time. However, in fact we found that (1) both functions decrease over time (2) with species-rich plant communities being less affected than species-poor plant communities (predation), (3) both functions are positively affected by increasing plant species richness ^see^ ^also^ ^70,71^, and (4) the positive effect of plant species richness on predation, but not herbivory becomes stronger over time.

There are two possible explanations for this uncoupling of biomass ratios (stocks) and ecosystem functions (flows). First, methodological limitations might play a role. Our arthropod sampling does not fully capture all groups contributing to predation and herbivory. For example, we exposed the dummy caterpillars for 24 hours, whereas the arthropod sampling was conducted during the day only. That means, with the suction sampling we missed all night-active predatory arthropods, for example some carabids and staphylinids. Further, with the assessment of leaf damage during the time of maximal biomass production, we recorded accumulated damage by various herbivores occurring over the entire growing season. That means we also recorded damage caused by arthropod groups, which are not present during the time of suction sampling or that are not caught at all by our regular sampling (e.g. slugs). Second, changes in ecosystem functions (herbivory and predation) may be driven less by the quantity of arthropods and more by their functional composition. As ecosystems become more diverse, shifts in traits such as food specialization, feeding mode and overall trait diversity ^34^ might increase the functioning of arthropod communities ^72–74^. In addition, higher functional complementarity, both in space and time, could lead to more efficient resource use, even in environments with stable or declining arthropod biomass ^34,54,75,76^. However, comprehensive trait-based analyses would be needed to support the proposed mechanism.

Overall, our results underline that the main and interactive effects of time and plant species richness on ecosystem functions like herbivory and predation are not simply a result of biomass ratios. Analysing compositional shifts in arthropod communities, but also applying a more complete arthropod sampling at a higher temporal resolution would be crucial to understand temporal variation in the plant species richness-ecosystem functioning relationship.

## Conclusion

Our findings confirm that diversification in semi-natural grasslands immediately and consistently supports arthropod communities. Especially in summer, when environmental conditions are harsh, fewer species are lost in high-diversity compared to low-diversity grasslands, so we suggest that high plant species richness in semi-natural grasslands has the potential to stabilize arthropod communities over time. However, this stabilizing effect of plant diversity may take years to become fully established, emphasizing the value of long-term conservation efforts. Since temporal variation in plant species richness effects cannot be fully explained by plant productivity alone, future research should explore the roles of changing abiotic factors and successional dynamics in grasslands. Likewise shifts in arthropod biomass and biomass ratios among trophic levels could not explain the temporal dynamics in arthropod community function over time underlining the need for more detailed studies of shifts in arthropod community and trait composition to better understand the mechanisms behind these patterns.

## Methods

### The Jena Experiment

The Jena Experiment was established in 2002 in the floodplain of the river Saale (Thuringia, Germany, 50°550 N, 11°350 E) and uses 60 plant species native to Central European mesophilic grasslands. Plant communities were sown in 82 plots of 20 x 20 m with a gradient of species richness (1, 2, 4, 8, 16 and 60) arranged in four spatial blocks ^77^. This setup includes 16 replicates each for monocultures, 2-species, 4-species, and 8-species mixtures, along with 14 replicates for 16-species and 4 replicates for 60-species mixtures (totaling 82 plots). In 2009, plot sizes were reduced to 100 square meters, and monocultures of *Bellis perennis* (L., 1753) and *Cynosurus cristatus* (L., 1753) were abandoned due to insufficient cover of the target species, leaving 80 plots for the current study ^30^. The specific species mixtures were randomly chosen under certain constraints ^77^. Regular maintenance involves weeding twice per year, which sustains the plant species richness gradient and maintains a strong correlation between the sown and realized numbers of target species ^30^. Following the mowing regime typical for extensively managed hay meadows in Central Germany, the plots are mown at the peak biomass period in early June and early September, with the hay being removed afterward.

### Measuring plant biomass

We measured aboveground plant biomass in every year between 2010 and 2022 ^31^. Always at the time of peak biomass in late May and August, we harvested plant material in two frames of 0.1 m² each on every single plot. Finally, we sorted the samples by species, dried them for 72 hours at 70°C and weighed them. To quantify community productivity, we summed up dried biomass of all target species per sample, averaged values of both samples and extrapolated to 1 m².

### Measuring plant species cover

Within an area of 3 m x 3 m we measured the species-specific plant cover in every plot in every year during the time of peak biomass (late May and August). We estimated cover of each target species, and the total cover of all target species using the Londo scale (Londo 1976).

### Estimation of herbivory

Between 2010 and 2022, we measured leaf damage on all species sown on the respective plot once per year in late August (second biomass harvest, see above) ^48,71^. Data collection resulted in leaf damage data for 3,020 plant species x plot x year combinations. For each species, we assessed damage on 20 randomly selected leaves per plant species (or all leaves if fewer than 20 leaves were available) from the biomass sample (see section: Measuring plant biomass). First, we visually estimated the absolute area damaged by (1) chewing, (2) rasping, and (3) leaf-mining herbivores (in mm²) using a template card containing various circles and squares of known sizes for reference. Second, we used a leaf area meter (LI-3000C Area Meter, LI-COR Biosciences, Lincoln, Nebraska, USA) to measure the leaf area of all leaves per species. As the area meter solely calculates the remaining area, encompassing rasping and mining damage but excluding chewing damage, we estimated the initial leaf area by adding the area lost due to chewing damage. For our analysis, we calculated the percentage herbivory of each species in a plot by dividing the damaged leaf area by the initial leaf area and multiplying by 100. Plot-level herbivory was calculated as the abundance-weighted mean of species-specific herbivory percentages (weighted by leaf biomass for each plot). Overall, the species for which we estimated herbivory were representative of the composition of the plant communities in the plots. On median, they covered more than 80% of the plant community growing within a 3×3 m area of the plot (see Figure S12). This coverage did not change significantly over time or with changes in plant diversity.

### Estimation of predation

Between 2014 and 2022, we measured predation rates by invertebrates once per year and plot by using artificial dummies ^70^. The cylinder-shaped dummies (0.6 x 2.5 cm in size) were made from green plasticine (Staedtler Noris Club, Nuremberg, Germany) and mimic lepidopteran larvae ^70,78^. Always at peak biomass in late August, we placed 10 dummies per plot on the ground with a distance of 20 cm from each other. After 24 hours, we inspected the dummies for mandibular marks, stylet (predatory bugs) and ovipositor marks (e.g., from parasitic Hymenoptera). We recorded binary data for the presence or absence of any of these bite marks. The percentage of attacked dummies served as a measure of predation rate per plot and year.

### Sampling of arthropods and their functional traits

Between 2010 and 2016, we sampled aboveground arthropods every two years, and from 2017 onwards (until 2020) every year. We collected arthropods by suction sampling on all 80 plots twice per year, in May (spring) and July (summer; only May sampling in 2018). Using a modified commercial vacuum cleaner (Kärcher A2500; Kärcher GmbH, Winnenden, Germany), we randomly selected two subplots of 0.75 m x 0.75 m within each plot, covered them with a gauze-coated cage of the same size, and sampled until no more arthropods were visible in the cage. We sampled between 9 am and 4 pm on sunny days without wind. In May 2019 we only sampled one replicate per plot, due to unfavorable weather conditions. All species groups that can be representatively sampled with our method (Araneae, Coleoptera, Hemiptera, Hymenoptera) were sent to specialized taxonomists for identification. Due to difficulties in identification caused by larval stages, small sizes (e.g., hymenopterans), or damaged individuals, we could not identify all individuals to the species level. For our analyses, we used the highest taxonomic resolution that was reliably identifiable (71% species, 10% genus, 9% subfamily, and 10% family level). For all taxa that were sampled more than once in suction campaigns (non-singletons), we compiled information about feeding guild and wet body mass ^79^. We followed the published definition and divided taxa into herbivores (primarily feeding on plant material), predators (including parasitoids; primarily feeding on animals), detritivores (primarily feeding on dead organic matter) and omnivores (consume more than one resource type). For each year, sampling campaign and plot, we first calculated species richness, extrapolated species richness (Chao estimate ^80^), abundance, Hill number ^81^, biomass (mg/m²) and the biomass ratio (arthropod biomass in mg/m² divided by plant biomass in g/m²) for each sample within a plot and then averaged the two values per plot. We repeated the calculation for the subset of herbivores (biomass ratio here: herbivore biomass/ plant biomass) and predators (biomass ratio here: predator biomass/ herbivore biomass). We generated plant biomass from a 0.4 m x 0.5 m, however, this area covers a large proportion of the species found in a 3 x 3 m area during cover estimation. This means that the heterogeneity of the plant community in a plot is well covered, and extrapolating the plant biomass to the size of the insect cage (0.75 m x 0.75 m) is justifiable (see Figures S12). We calculated a total of eighteen metrics describing arthropod communities, for each combination of plot x sampling x year x season.

### Statistical analyses

All data processing and analyses were conducted using the statistical program R, version 4.3.3 ^82^.

### Models testing for temporal trends and its dependence on plant species richness

With a first mixed-effects model (Type III Sums of Squares; ‘lmer’-function, ‘lme4’ package ^83^; ‘lmerTest’ package ^84^) we tested for overall changes in the response variable over **time** (calendar year fitted as a centered numerical variable), in both sampling **seasons** (categorical variable with two levels) across a gradient in plant species richness (**PSR**; log transformed and centered). For herbivory and predation, season was not fitted in the model, as only one sampling campaign is available. Degrees of freedom were calculated using the Satterthwaite method.

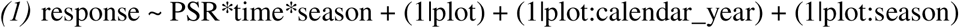

All models contained plot identity as a random term to account for spatial non-independence of the data. Moreover, if the response variable was measured multiple times in the same year the combination of plot and calendar year (nine levels) and the interaction of plot and season (two levels) were fitted as random effects to account for spatio-temporal non-independence of the data. Further, we use the model to estimate the relative annual changes in the response (fitted values for year i / fitted values for year i+1) in high (60 plant species) and low (monoculture) species mixtures.

### Models testing for plant species richness effects across multiple years

In a second mixed-effects model, we tested if **PSR** affects the response variable in different **calendar years** (fitted as a factor) and, if available, sampling **season** (fitted as a two-level factor).

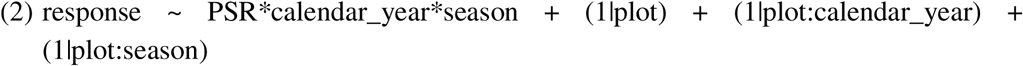

Models contained the same random terms as above. We use this model to extract the PSR effect on the response variable for every year (slope of the interaction of PSR and calendar year, or the interaction of PSR, calendar year and season). Each slope was divided by the average (across all plots) of the respective year or season within a year to make slopes independent of absolute differences between years and seasons (**corrected PSR effect** on the response variable).

### Models testing for temporal changes in the effects of plant species richness

Lastly, in a linear model we tested for directional changes in plant species richness (PSR) effects (slopes) over **time** (calendar year fitted as centered, numerical variable) for both **seasons**.

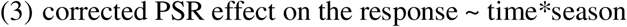

## Acknowledgements

We thank the technical staff of the Jena Experiment for their work in maintaining the experimental field site and also many student helpers for weeding of the experimental plots and support during measurements. Further, we thank the speaker and central coordination team of the Jena Experiment, Nico Eisenhauer, Alexandra Weigelt, Christiane Roscher and Gerd Gleixner, for guiding the Research Unit and providing plant cover and biomass data. Roland Achtziger, Eric Anton, Theo Blick, Frank Creutzburg, Ralf Heckmann, Christoph Muster, and Oliver Wiche is acknowledged for their tremendous work with identifying arthropod individuals and Alex Strauss and two anonymous reviewers for helpful comments on the manuscript. This work was supported by the German Science Foundation (DFG FOR 1451 EB 555/3-1, WE 3081/15-2, FOR 5000 EB 555/6-1, EB 555/6-2, ME 5474/1-1, ME 5474/1-2).

## Author contributions

A.E., and S.T.E. developed and framed the research question. A.E. and M.B. analysed the data. H.S. and S.T.M. contributed to the data analysis. A.E. wrote the manuscript with contributions from all other authors. A.E., L.H. and M.B. contributed to data collection.

## Data accessibility statement

Data supporting the results of this study are available from the Jena Experiment Information System (JEXIS) at https://doi.org/10.25829/V1ND-PG20.

