## Supplemental Tables 1-7; Supplemental Figures 1-12 for "Plant diversity promotes aboveground arthropods and associated functions despite arthropod loss over time"

#### ^2^Laboratoire d’Inventaire Forestier, Ecole Nationale des Sciences Géographiques-Géomatique, Institut National de l'Information Géographique et Forestière, Nancy, France)

#### ^3^Terrestrial Ecology Research Group, School of Life Sciences, Technical University of Munich, Freising, Germany

^†^ these authors contributed equally

**Table S1: Temporal trends in arthropod community metrics and its dependence on plant species richness (model 1).** Results of linear mixed-effects models relating species richness, estimated richness (Chao estimate), abundance, Hill number and biomass (mg/m²) of total arthropods, herbivores and predators to the explanatory variables plant species richness (PSR), time (calendar year as numerical variable) sampling season (spring/ May and summer/ July) and all possible interactions between PSR, time and season. For a detailed model description see model (1) in the section *Statistical analysis*. For each explanatory variable the F-statistics and degree of freedom are shown. Abundance and biomass were log-transformed prior to the analyses. Significances are indicated by asterisks as followed: *** *P* < 0.001; ** *P* < 0.01; * *P* < 0.05; (*) *P* < 0.1.

|  | **Richness** | **Estimated richness** | **Abundance** | **Hill number** | **Biomass** |
| --- | --- | --- | --- | --- | --- |
| **Total** | | | | | |
| Plant species richness | F₁,₈₀ = 133.5*** | F₁,₁₅₇ = 111.4*** | F₁,₈₀ = 100.7*** | F₁,₈₁ = 81.1*** | F₁,₈₁ = 80.6*** |
| Time | F₁,₅₄₄ = 224.5*** | F₁,₁₀₁₅ = 148*** | F₁,₅₂₅ = 231.4*** | F₁,₅₃₅ = 128.7*** | F₁,₅₁₆ = 156.1*** |
| Season | F₁,₈₀ = 321.8*** | F₁,₁₅₇ = 127.9*** | F₁,₈₁ = 538.6*** | F₁,₈₁ = 136.2*** | F₁,₈₁ = 209.5*** |
| PSR x Time | F₁,₅₄₃ = 10.8** | F₁,₁₀₁₅ = 4.8* | F₁,₅₂₅ = 1.3 | F₁,₅₃₅ = 3.2(*) | F₁,₅₁₆ = 0.1 |
| PSR x Season | F₁,₈₀ = 43.6*** | F₁,₁₅₇ = 20.0*** | F₁,₈₁ = 22.2*** | F₁,₈₁ = 15.6*** | F₁,₈₁ = 7.5** |
| Time x Season | F₁,₅₄₆ = 1.5 | F₁,₁₀₁₅ = 3.6(*) | F₁,₅₂₈ = 4.0* | F₁,₅₃₃ = 22.6*** | F₁,₅₁₃ = 11.5*** |
| PSR x Time x Season | F₁,₅₄₅ = 0.5 | F₁,₁₀₁₅ = 1.1 | F₁,₅₂₈ = 1.9 | F₁,₅₃₃ = 4.1* | F₁,₅₁₃ = 1.0 |
| **Herbivores** | | | | | |
| Plant species richness | F₁,₈₁ = 140.2*** | F₁,₈₀ = 120.2*** | F₁,₈₀ = 70.7*** | F₁,₈₀ = 69.1*** | F₁,₈₁ = 82.1*** |
| Time | F₁,₅₃₈ = 109.1*** | F₁,₁₀₄₃ = 39.9*** | F₁,₁₀₂₆ = 155.5*** | F₁,₄₉₉ = 40.6*** | F₁,₅₁₁ = 113.5*** |
| Season | F₁,₈₁ = 573.1*** | F₁,₁₀₄₇ = 199*** | F₁,₈₁ = 670.3*** | F₁,₈₃ = 234.1*** | F₁,₆₀₃ = 450.9*** |
| PSR x Time | F₁,₅₃₈ = 3.0(*) | F₁,₁₀₄₂ = 2.7 | F₁,₁₀₂₆ = 0.7 | F₁,₄₉₇ = 0.6 | F₁,₅₁₁ = 0.1 |
| PSR x Season | F₁,₈₁ = 60.9*** | F₁,₁₀₄₈ = 32.0*** | F₁,₈₁ = 28.7*** | F₁,₈₂ = 22.6*** | F₁,₆₀₃ = 18.3*** |
| Time x Season | F₁,₅₄₂ = 0.3 | F₁,₁₀₄₃ = 4.3* | F₁,₁₀₂₆ < 0.1 | F₁,₅₀₂ = 9.2** | F₁,₅₈₄ = 9.9** |
| PSR x Time x Season | F₁,₅₄₂ = 4.7* | F₁,₁₀₄₂ = 15.8*** | F₁,₁₀₂₆ = 3.1(*) | F₁,₅₀₀ = 9.5** | F₁,₅₈₄ = 3.2(*) |
| **Predators** | | | | | |
| Plant species richness | F₁,₈₁ = 75.1*** | F₁,₈₃ = 50*** | F₁,₈₁ = 85.4*** | F₁,₈₅ = 64.2*** | F₁,₈₁ = 59.5*** |
| Time | F₁,₅₄₉ = 194.4*** | F₁,₅₃₆ = 154.2*** | F₁,₅₃₃ = 195.8*** | F₁,₅₄₃ = 157.1*** | F₁,₅₂₃ = 154.2*** |
| Season | F₁,₈₀ = 95.3*** | F₁,₈₂ = 41.6*** | F₁,₈₀ = 108*** | F₁,₈₃ = 65.7*** | F₁,₆₂₈ = 49.4*** |
| PSR x Time | F₁,₅₄₉ = 9.4** | F₁,₅₄₄ = 2.1 | F₁,₅₃₂ = 4.3* | F₁,₅₅₀ = 5.0* | F₁,₅₂₃ = 0.5 |
| PSR x Season | F₁,₈₀ = 19.5*** | F₁,₈₄ = 8.1** | F₁,₈₀ = 11.5** | F₁,₈₅ = 13.4*** | F₁,₆₂₈ = 3.0(*) |
| Time x Season | F₁,₅₄₅ = 4.6* | F₁,₅₃₁ = 0.6 | F₁,₅₃₆ = 19.5*** | F₁,₅₃₃ = 3.5(*) | F₁,₆₀₈ = 9.9** |
| PSR x Time x Season | F₁,₅₄₅ = 0.2 | F₁,₅₃₉ = 3.2(*) | F₁,₅₃₅ = 0.4 | F₁,₅₄₂ = 0.1 | F₁,₆₀₈ = 0.1 |

**Table S2:** Annual (%) change in arthropod community metrics in plant communities of low species richness (monocultures), intermediate species richness (8 plant species) and high species richness (60 plant species). Values are predicted changes extracted from model (1), testing for overall changes in the response variable over time, in both sampling seasons across a gradient in plant species richness. Given are mean values (average loss across both sampling seasons), and values for spring and summer samplings, respectively.

|  | **PSR level** | **Annual change Mean (%)** | **Annual change Spring (%)** | **Annual change Summer (%)** |
| --- | --- | --- | --- | --- |
| **Total** | | | | |
| **Richness** | low | -5.3 | -6.3 | -4.5 |
|  | intermediate | -5.2 | -6.9 | -4.1 |
|  | high | -5.2 | -7.5 | -3.7 |
| **Estimated richness** | low | -5.3 | -6.2 | -4.6 |
| **(Chao estimate)** | intermediate | -5.1 | -6.7 | -3.8 |
|  | high | -4.9 | -7.1 | -3.2 |
| **Abundance** | low | -6.4 | -6.7 | -6.2 |
|  | intermediate | -6.7 | -7.3 | -6.0 |
|  | high | -7.1 | -8.2 | -5.8 |
| **Hill number** | low | -4.4 | -5.6 | -3.4 |
|  | intermediate | -4.2 | -6.2 | -2.5 |
|  | high | -4.1 | -6.7 | -1.8 |
| **Biomass** | low | -7.5 | -8.0 | -7.0 |
| **(mg/m²)** | intermediate | -7.4 | -8.2 | -6.4 |
|  | high | -7.2 | -8.4 | -5.2 |
| **Herbivores** |  |  |  |  |
| **Richness** | low | -4.2 | -4.3 | -4.2 |
|  | intermediate | -3.9 | -5.9 | -2.8 |
|  | high | -3.7 | -7.1 | -1.8 |
| **Estimated richness** | low | -3.1 | -0.7 | -4.4 |
| **(Chao estimate)** | intermediate | -3.2 | -5.3 | -1.8 |
|  | high | -3.3 | -7.9 | +0.4 |
| **Abundance** | low | -6.0 | -5.6 | -6.5 |
|  | intermediate | -6.1 | -6.7 | -6.0 |
|  | high | -6.5 | -7.9 | -5.2 |
| **Hill number** | low | -2.5 | -2.3 | -2.5 |
|  | intermediate | -2.3 | -4.0 | -1.1 |
|  | high | -2.3 | -5.6 | +0.5 |
| **Biomass** | low | -7.2 | -7.6 | -7.0 |
| **(mg/m²)** | intermediate | -7.0 | -8.0 | -5.7 |
|  | high | -6.6 | -8.5 | -2.8 |
| **Predators** |  |  |  |  |
| **Richness** | low | -6.2 | -7.6 | -4.8 |
|  | intermediate | -6.1 | -7.6 | -4.9 |
|  | high | -6.0 | -7.5 | -5.0 |
| **Estimated richness** | low | -6.7 | -8.2 | -5.0 |
| **(Chao estimate)** | intermediate | -6.1 | -7.0 | -5.2 |
|  | high | -5.5 | -5.7 | -5.4 |
| **Abundance** | low | -6.3 | -7.4 | -4.9 |
|  | intermediate | -6.8 | -7.9 | -5.4 |
|  | high | -7.5 | -8.5 | -6.1 |
| **Hill number** | low | -5.0 | -6.0 | -4.0 |
|  | intermediate | -5.0 | -6.1 | -4.1 |
|  | high | -5.0 | -6.1 | -4.2 |
| **Biomass** | low | -7.9 | -8.7 | -6.6 |
| **(mg/m²)** | intermediate | -8.1 | -8.7 | -7.1 |
|  | high | -8.4 | -8.9 | -7.8 |

**Table S3:** Annual (%) change in the biomass ratios of arthropods and plants (total mg arthropod biomass to total g plant biomass), herbivores (mg) and plants (g), and predators (mg) and herbivores (mg), percentage herbivory and predation rate in plant communities of low species richness (monocultures), intermediate species richness (8 plant species) and high species richness (60 plant species). Values are predicted changes extracted from model (1), testing for overall changes in the response variable over time, in both sampling seasons (for biomass ratios only) across a gradient in plant species richness. Given are mean values (average loss across both seasons), and values for spring and summer samplings, respectively.

|  | **PSR level** | **Annual change Mean (%)** | **Annual change Spring (%)** | **Annual change Summer (%)** |
| --- | --- | --- | --- | --- |
| **Arthropod load** | low | +6.5 | +2.2 | +11.6 |
|  | intermediate | +0.4 | -3.2 | +4.2 |
|  | high | -7.8 | -14.1 | -4.1 |
| **Herbivore-plant** | low | +11.9 | +9.9 | +14.0 |
| **ratio** | intermediate | +4.6 | +1.2 | +7.6 |
|  | high | -7.3 | -24.2 | -1.1 |
| **Predator-herbivore** | low | -0.7 | -1.5 | +0.5 |
| **ratio** | intermediate | -1.3 | -0.6 | -2.1 |
|  | high | -2.2 | -0.9 | -5.8 |
| **Herbivory** | low | -4.2 |  |  |
|  | intermediate | -4.8 |  |  |
|  | high | -5.5 |  |  |
| **Predation** | low | -11.9 |  |  |
|  | intermediate | -9.1 |  |  |
|  | high | -5.4 |  |  |

**Table S4: Temporal trends in herbivory, predation and biomass ratios and its dependence on plant species richness (model 1).** Results of linear mixed-effects models relating the biomass ratios of arthropods (mg) and plants (g), herbivores (mg) and plants (g), and predators (mg) to herbivores (mg), percentage herbivory and predation rate to the explanatory variables plant species richness (PSR), time (calendar year as numeric variable), sampling seasons (spring/ May and summer/ July; for biomass ratios only) and all possible interactions between PSR, time and season. For a detailed model description see model (1) in the section *Statistical analysis*. Arthropod biomass ratios and herbivory were log transformed prior to the analyses. Significances are indicated by asterisks as followed: *** *P* < 0.001; ** *P* < 0.01; * *P* < 0.05; (*) *P* < 0.1.

|  | **Arthropod-plant ratio** | **Herbivore-plant ratio** | **Predator-herbivore ratio** | **Herbivory** | **Predation** |
| --- | --- | --- | --- | --- | --- |
| PSR | F₁,₇₅ = 29.4*** | F₁,₇₄ = 27.5*** | F₁,₈₂ = 0.7 | F₁,₈₂ = 4.3* | F₁,₈₀ = 38.9*** |
| Time | F₁,₄₇₆ = 0.1 | F₁,₄₄₄ = 7.7** | F₁,₄₆₇ = 1.4 | F₁,₈₂₂ = 67.1*** | F₁,₆₄₈₈ = 340.9*** |
| Season | F₁,₇₄ = 104*** | F₁,₅₃₂ = 104.1*** | F₁,₅₈₈ = 132.9*** |  |  |
| PSR x Time | F₁,₄₇₇ = 10.7** | F₁,₄₄₄ = 9.1** | F₁,₄₇₃ = 0.2 | F₁,₈₂₆ = 2.6 | F₁,₆₄₈₈ = 5.9* |
| PSR x Season | F₁,₇₆ = 9.5** | F₁,₅₂₀ = 12.5*** | F₁,₅₈₈ = 3.6(*) |  |  |
| Time x Season | F₁,₄₇₉ = 11.2*** | F₁,₄₉₄ = 6.4* | F₁,₅₄₂ = 0.3 |  |  |
| PSR x Time x Season | F₁,₄₈₀ < 0.1 | F₁,₄₉₀ = 0.5 | F₁,₅₄₈ = 1.2 |  |  |

**Table S5: Plant species richness effects on arthropod community metrics in multiple years (model 2).** Results of linear mixed-effects models relating species richness, estimated richness (Chao estimate), abundance, Hill number and biomass (mg/m²) of the total arthropod communities, herbivores and predators to the explanatory variables plant species richness (PSR), calendar year (CY, as factor), sampling season (spring/ May and summer/ July) and all possible interactions between PSR, calendar year and season. For a detailed model description see model (2) in the section *Statistical analysis*. Abundance and biomass were log-transformed prior to the analyses. Significances are indicated by asterisks as followed: *** *P* < 0.001; ** *P* < 0.01; * *P* < 0.05; (*) *P* < 0.1.

|  | **Richness** | **Estimated richness** | **Abundance** | **Hill number** | **Biomass** |
| --- | --- | --- | --- | --- | --- |
| **Explanatory variable** | | | | | |
| **Total** |  |  |  |  |  |
| Plant species richness | F₁,₈₁ = 130.0*** | F₁,₁₆₀ = 108.1*** | F₁,₈₁ = 100.2*** | F₁,₈₂ = 77.2*** | F₁,₈₁ = 82.7*** |
| Calendar year | F₇,₅₆₆ = 60.7*** | F₇,₁₀₁₈ = 36.3*** | F₇,₅₄₆ = 62.3*** | F₇,₅₅₈ = 58.0*** | F₇,₅₃₂ = 52.9*** |
| Season | F₁,₈₇ = 286.9*** | F₁,₁₇₃ = 114.4*** | F₁,₈₇ = 480.8*** | F₁,₈₈ = 120.7*** | F₁,₈₈ = 185.1*** |
| PSR x CY | F₇,₅₆₆ = 3.1** | F₇,₁₀₁₉ = 1.7 | F₇,₅₄₅ = 2.8** | F₇,₅₅₉ = 2.1* | F₇,₅₃₂ = 1.5 |
| PSR x Season | F₁,₈₇ = 43.6*** | F₁,₁₇₃ = 20.7*** | F₁,₈₇ = 24.8*** | F₁,₈₈ = 16.2*** | F₁,₈₈ = 10.4** |
| CY x Season | F₆,₅₂₈ = 15.1*** | F₆,₁₀₁₈ = 13.3*** | F₆,₅₀₁ = 30.5*** | F₆,₅₂₁ = 22.9*** | F₆,₄₈₈ = 20.8*** |
| PSR x CY x Season | F₆,₅₂₇ = 2.4* | F₆,₁₀₁₈ = 2.7* | F₆,₅₀₁ = 2.3* | F₆,₅₂₂ = 3.4** | F₆,₄₈₈ = 2.7* |
| **Herbivores** |  |  |  |  |  |
| Plant species richness | F₁,₈₂ = 134.0*** | F₁,₈₇ = 113.7*** | F₁,₈₁ = 68.3*** | F₁,₈₅ = 67.3*** | F₁,₈₂ = 81.3*** |
| Calendar year | F₇,₅₆₆ = 34.5*** | F₇,₉₇₈ = 11.8*** | F₇,₅₄₉ = 50.3*** | F₇,₅₅₃ = 19.5*** | F₇,₅₂₅ = 57.2*** |
| Season | F₁,₈₈ = 503.4*** | F₁,₉₁ = 195.1*** | F₁,₈₇ = 590.4*** | F₁,₉₃ = 223.0*** | F₁,₈₉ = 392.3*** |
| PSR x CY | F₇,₅₆₅ = 1.8(*) | F₇,₉₇₆ = 2.3* | F₇,₅₄₉ = 1.6 | F₇,₅₄₇ = 0.7 | F₇,₅₂₅ = 0.9 |
| PSR x Season | F₁,₈₈ = 58.6*** | F₁,₈₈ = 31.0*** | F₁,₈₇ = 29.0*** | F₁,₉₀ = 22.1*** | F₁,₈₉ = 19.9*** |
| CY x Season | F₆,₅₂₆ = 16.9*** | F₆,₉₇₃ = 7.6*** | F₆,₅₀₉ = 27.9*** | F₆,₅₁₃ = 11.7*** | F₆,₄₈₆ = 22.3*** |
| PSR x CY x Season | F₆,₅₂₅ = 2.6* | F₆,₉₆₉ = 6.1*** | F₆,₅₀₉ = 1.7 | F₆,₅₀₆ = 4.1*** | F₆,₄₈₆ = 1.7 |
| **Predators** |  |  |  |  |  |
| Plant species richness | F₁,₈₁ = 73.7*** | F₁,₈₄ = 50.9*** | F₁,₈₁ = 84.1*** | F₁,₈₅ = 64.0*** | F₁,₈₁ = 61.4*** |
| Calendar year | F₇,₅₆₇ = 69.2*** | F₇,₅₆₀ = 41.2*** | F₇,₅₄₈ = 68.6*** | F₇,₅₆₂ = 53.2*** | F₇,₅₄₄ = 40.1*** |
| Season | F₁,₈₈ = 86.4*** | F₁,₉₀ = 37.6*** | F₁,₈₈ = 100.9*** | F₁,₉₁ = 60.5*** | F₁,₉₁ = 50.6*** |
| PSR x CY | F₇,₅₆₆ = 3.9*** | F₇,₅₇₁ = 1.0 | F₇,₅₄₈ = 3.3** | F₇,₅₇₄ = 2.1* | F₇,₅₄₄ = 1.3 |
| PSR x Season | F₁,₈₇ = 20.5*** | F₁,₉₂ = 8.2** | F₁,₈₈ = 13.5*** | F₁,₉₄ = 14.0*** | F₁,₉₁ = 5.0* |
| CY x Season | F₆,₅₂₇ = 14.8*** | F₆,₅₁₇ = 11.1*** | F₆,₅₁₆ = 18.4*** | F₆,₅₁₉ = 13.4*** | F₆,₅₀₉ = 12.2*** |
| PSR x CY x Season | F₆,₅₂₇ = 3.5** | F₆,₅₂₈ = 1.9(*) | F₆,₅₁₅ = 2.8* | F₆,₅₃₀ = 3.0** | F₆,₅₀₉ = 2.0(*) |

**Table S6**: **Changes in plant species richness effects over time (model 3).** Results from linear models testing whether plant species richness effects on arthropod estimated richness (Chao estimate), Hill number, abundance and biomass ratios (PSR slope) change over time (calendar year as numerical variable), between sampling seasons (spring/ May and summer/ July) or if there is an interactive effect of both (time x season). Biomass ratios shown here are: total arthropod biomass (mg) to total plant biomass (g), herbivore biomass (mg) to total plant biomass (g), and predator biomass (mg) to herbivore biomass (mg). For a detailed model description see model (3) in the section *Statistical analysis*. Significances are indicated by asterisks as followed: *** *P* < 0.001; ** *P* < 0.01; * *P* < 0.05; (*) *P* < 0.1.

|  | **Estimated richness** | **Abundance** | **Hill number** | **Arthropod- plant ratio** | **Herbivore-plant ratio** | **Predator-herbivore ratio** |
| --- | --- | --- | --- | --- | --- | --- |
| **Explanatory variable** | | | | | | |
| **Total** |  | | | | | |
| Time | F₁,₁₁ < 0.1 | F₁,₁₁ < 0.1 | F₁,₁₁ < 0.1 | F₁,₁₁ = 12.3** | F₁,₁₁ = 4.1(*) | F₁,₁₁ = 0.3 |
| Season | F₁,₁₁ = 1.0 | F₁,₁₁ = 1.1 | F₁,₁₁ = 1.5 | F₁,₁₁ = 21*** | F₁,₁₁ = 14.7** | F₁,₁₁ = 2.8 |
| Time x Season | F₁,₁₁ = 0.4 | F₁,₁₁ = 0.5 | F₁,₁₁ = 0.9 | F₁,₁₁ = 1.1 | F₁,₁₁ = 0.5 | F₁,₁₁ = 1.5 |
| **Herbivores** |  | | | | | |
| Time | F₁,₁₁ = 0.7 | F₁,₁₁ < 0.1 | F₁,₁₁ = 0.2 |  |  |  |
| Season | F₁,₁₁ = 2.5 | F₁,₁₁ = 1.6 | F₁,₁₁ = 3.9(*) |  |  |  |
| Time x Season | F₁,₁₁ = 5.2* | F₁,₁₁ = 1 | F₁,₁₁ = 4.1(*) |  |  |  |
| **Predators** |  | | | | | |
| Time | F₁,₁₁ = 0.6 | F₁,₁₁ = 0.1 | F₁,₁₁ = 0.1 |  |  |  |
| Season | F₁,₁₁ = 0.5 | F₁,₁₁ = 0.9 | F₁,₁₁ = 2.9 |  |  |  |
| Time x Season | F₁,₁₁ = 1.4 | F₁,₁₁ = 0.1 | F₁,₁₁ < 0.1 |  |  |  |

**Table S7: Plant species richness effects on herbivory, predation and biomass ratios in multiple years (model 2).** Results of linear mixed-effects models relating the biomass ratios of arthropods (mg) and plants (g), herbivores and plants, predators and herbivores, percentage herbivory and predation rate to the explanatory variables plant species richness (PSR), calendar year (CY, factor), sampling season (spring/ May and summer/ July; for biomass ratios only) and all possible interactions between PSR, CY and season. For a detailed model description see model (2) in the section *Statistical analysis*. Biomass ratios and herbivory were log transformed prior to the analyses. Significances are indicated by asterisks as followed: *** *P* < 0.001; ** *P* < 0.01; * *P* < 0.05; (*) *P* < 0.1.

|  | **Arthropod- plant ratio** | **Herbivore-plant ratio** | **Predator-herbivore ratio** | **Herbivory** | **Predation** |
| --- | --- | --- | --- | --- | --- |
| **Explanatory variable** | | | | | |
| PSR | F₁,₇₆ = 27.7*** | F₁,₇₇ = 21.2*** | F₁,₈₆ = 0.9 | F₁,₈₁ = 4.5* | F₁,₈₂ = 37.7*** |
| Calendar year (factor) | F₇,₅₁₃ = 15.3*** | F₇,₄₉₉ = 17.2*** | F₇,₅₂₆ = 5.8*** | F₁₁,₈₁₆ = 29.6*** | F₈,₆₄₈₈ = 94.6*** |
| Season | F₁,₈₁ = 79.2*** | F₁,₈₄ = 99.1*** | F₁,₅₇₄ = 139.1*** |  | |
| PSR x CY | F₇,₅₁₅ = 5.3*** | F₇,₄₉₃ = 5.8*** | F₇,₅₂₈ = 2.8** | F₁₁,₈₁₈ = 1.9* | F₈,₆₄₈₇ = 1.9(*) |
| PSR x Season | F₁,₈₂ = 12.6*** | F₁,₈₁ = 13.6*** | F₁,₅₆₆ = 1.0 |  |  |
| CY x Season | F₆,₄₇₂ = 14.2*** | F₆,₄₄₇ = 8.0*** | F₆,₅₅₄ = 5.1*** |  |  |
| PSR x CY x Season | F₆,₄₇₄ = 4.7*** | F₆,₄₄₀ = 3.8*** | F₆,₅₅₆ = 1.9(*) |  |  |

**Figure S1:** **Temporal trends in arthropod estimated richness (Chao Index), abundance and Hill number.** Regression lines and 95% confidence intervals show the predicted relationship between time (in years) and estimated richness (A), abundance (B) and Hill number (C). Both were derived from model (1), where time was included as a numerical variable. Arthropods were sampled from 2010 to 2020, and are separated into herbivores (blue) and predators (orange). Line styles indicate changes in spring (dotted), in summer (dashed) and across seasons (solid).

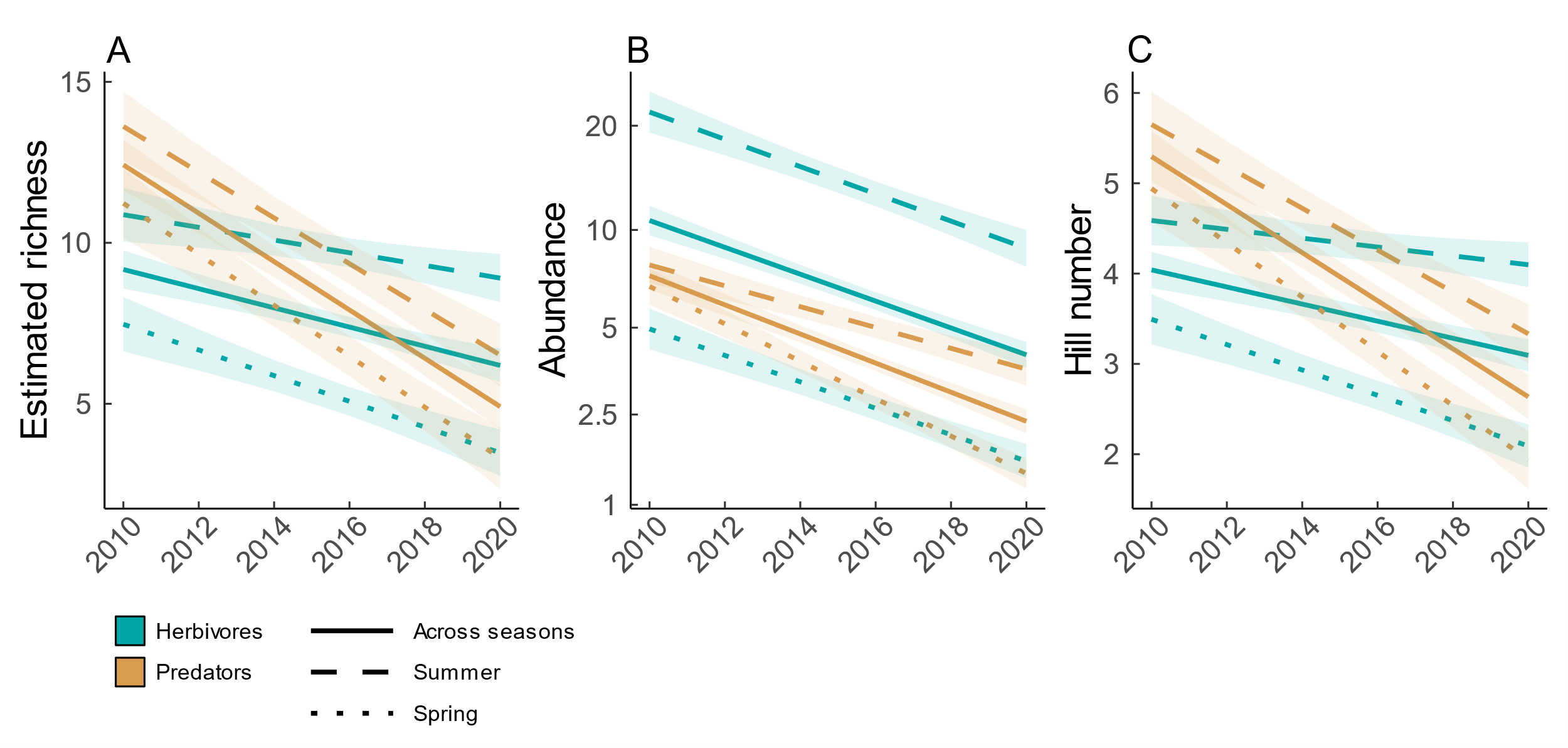

**Figure S2: Temporal trends in biomass ratios.** Regression lines and 95% confidence intervals show the predicted relationship between time (in years) and biomass ratios of arthropods and plants (A), herbivores and plants (B), and predators and herbivores (C). Both were derived from model (1), where time was included as a numerical variable. Arthropods and plants were sampled from 2010 to 2020. Biomass of arthropods was measured in mg/m² and biomass of plants in g/m². Line styles indicate changes in spring (dotted), in summer (dashed) and across seasons (solid).

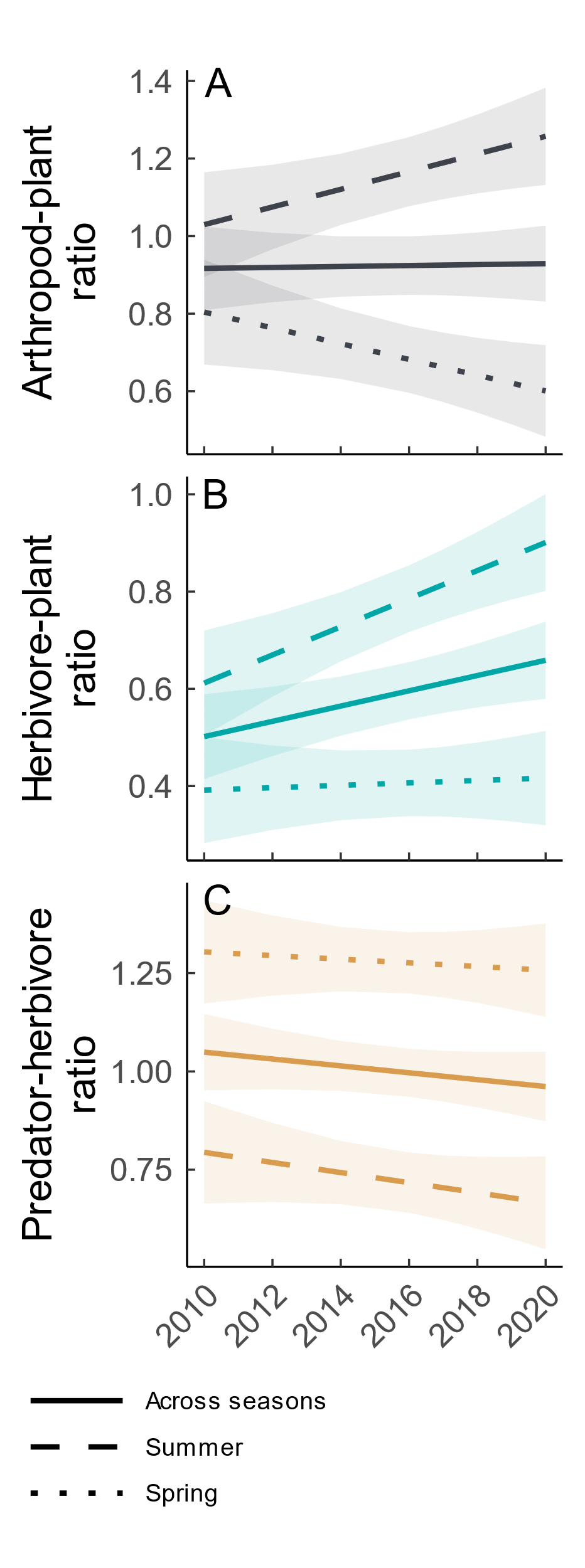

**Figure S3: Plant species richness effects on arthropod estimated richness (Chao estimate), abundance and Hill number in multiple years.** Regression lines show the predicted relationship between plant species richness and arthropod estimated richness (A-B), arthropod abundance (C-D), and arthropod Hill number (E-F) for each sampling year. Regression lines were derived from Model (2), where time was included as a factor, to allow for year-to year variation in the effects of plant species richness. Herbivores (blue) and predators (orange) were sampled between 2010 and 2020.

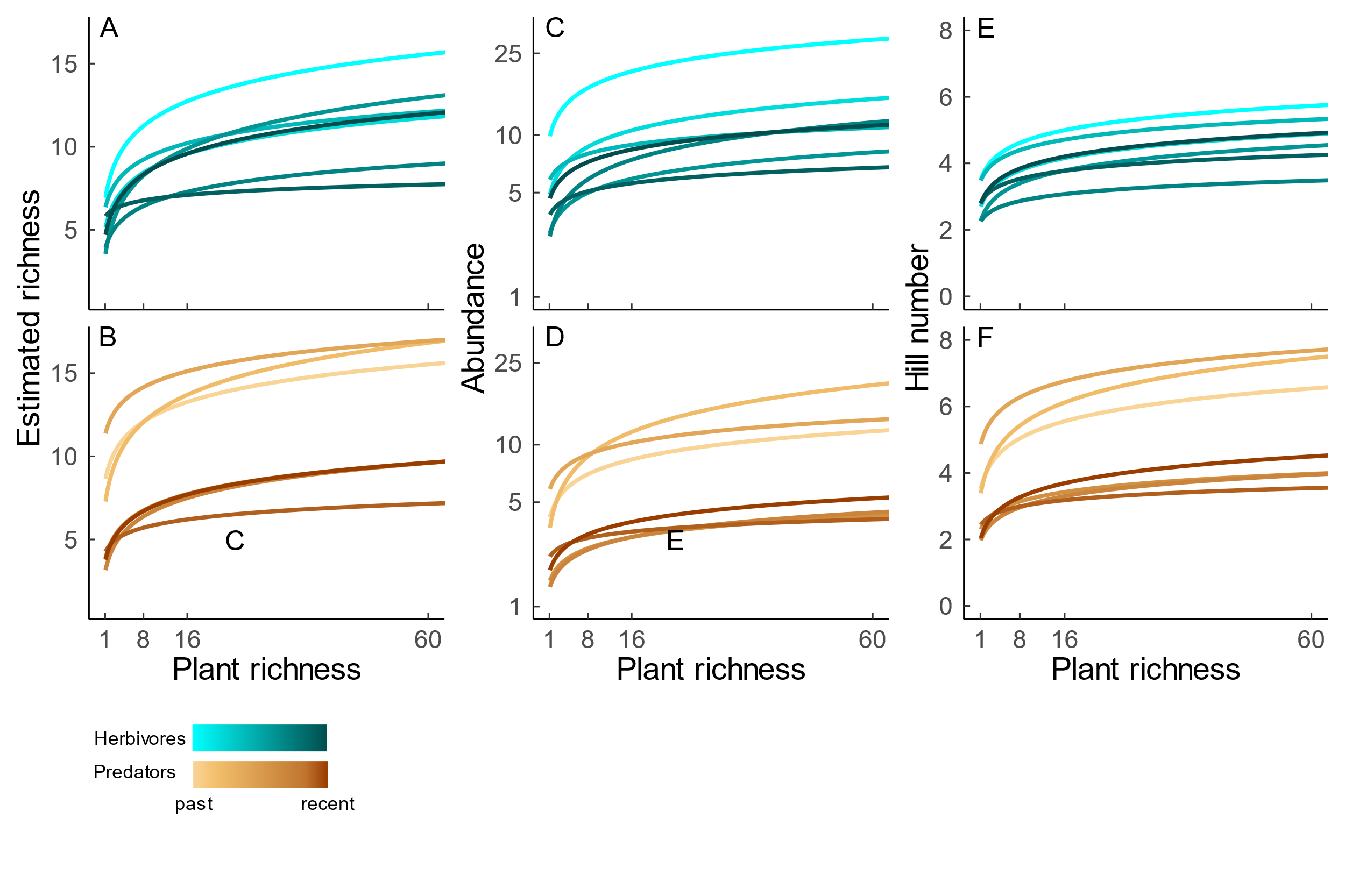

**Figure S4:** **Plant species richness effects on arthropod community metrics between 2010 and 2020.** Solid lines show main effects of plant species richness (plus confidence intervals) on total arthropod richness, estimated richness (Chao estimate), abundance, Hill number and biomass derived from model (2). Dotted and dashed lines show plant species richness effects for spring (May) and summer (July) sampling, respectively. In 2018 no main effects could be generated, as only arthropod data from May sampling was available.

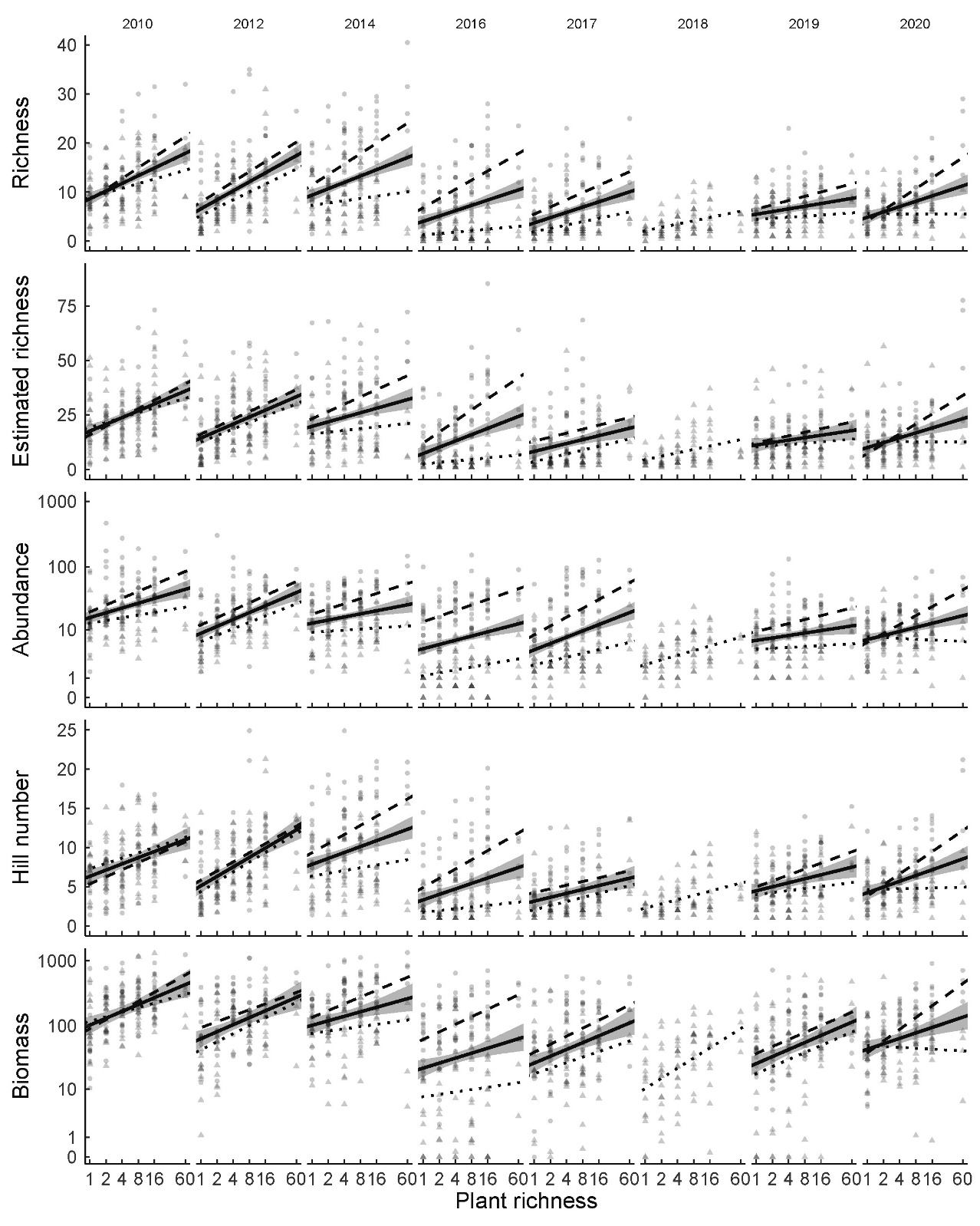

**Figure S5**: **Plant species richness effects on herbivore community metrics between 2010 and 2020.** Solid lines show main effects of plant species richness (plus confidence intervals) on herbivore richness, estimated richness (Chao estimate), abundance, Hill number and biomass derived from model (2). Dotted and dashed lines show plant species richness effects for spring (May) and summer (July) sampling, respectively. In 2018 no main effects could be generated, as only arthropod data from May sampling were available.

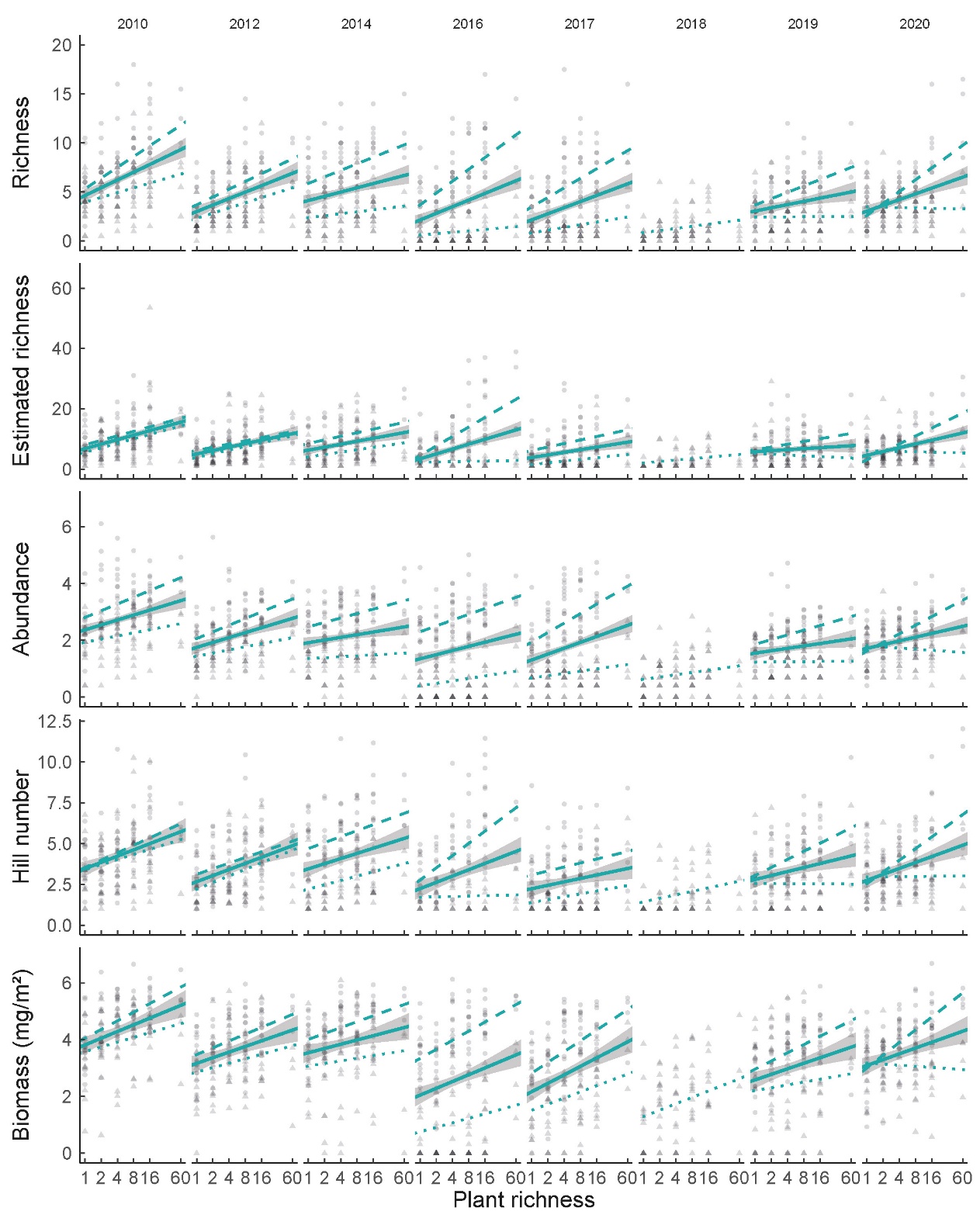

**Figure S6**: **Plant species richness effects on predator community metrics between 2010 and 2020.** Solid lines show main effects of plant species richness (plus confidence intervals) on predator richness, estimated richness (Chao estimate), abundance, Hill number and biomass derived from model (2). Dotted and dashed lines show plant species richness effects for spring (May) and summer (July) sampling, respectively. In 2018 no main effects could be generated, as only arthropod data from May sampling were available.
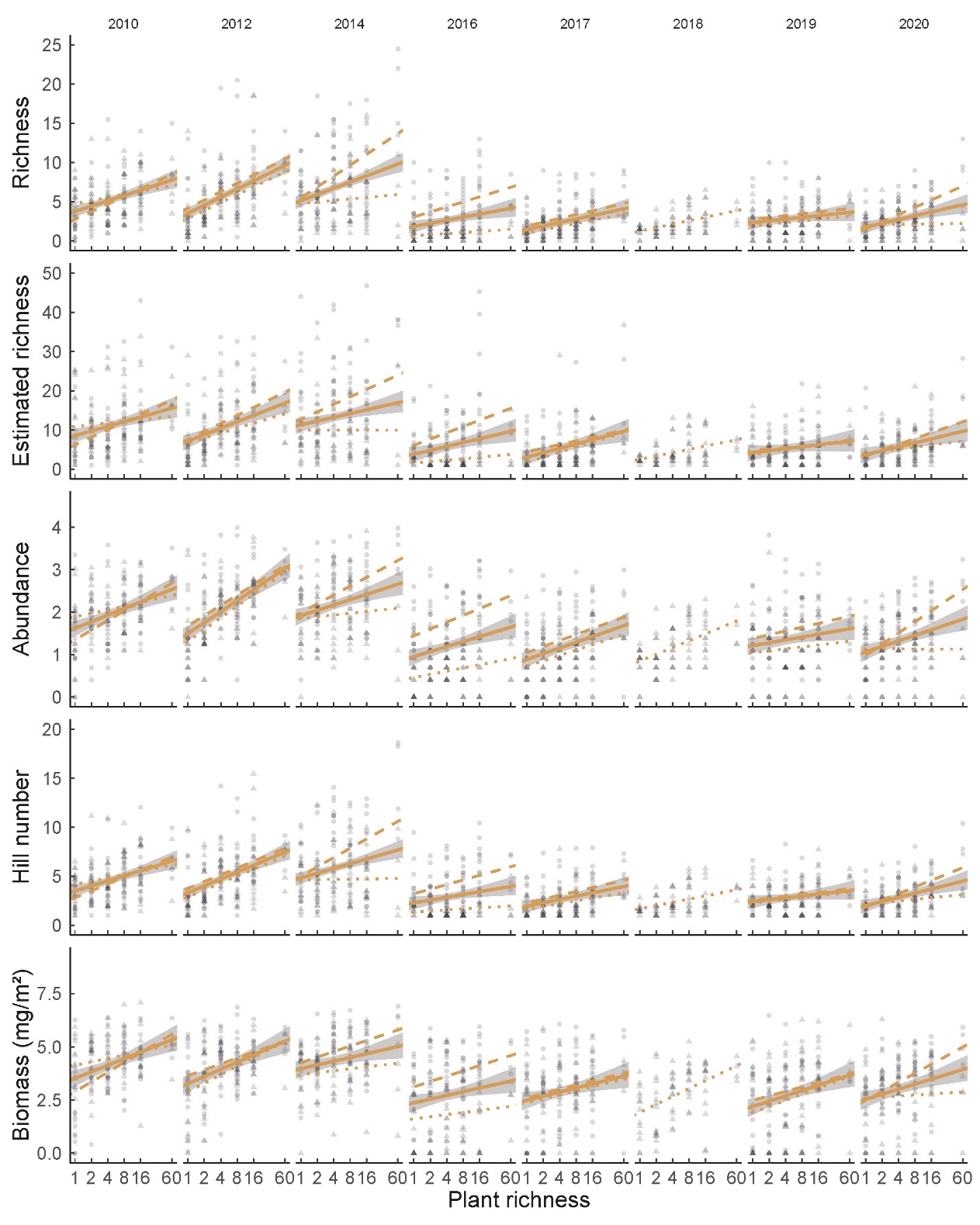

**Figure S7: Change in the effect of plant species richness on arthropod estimated richness, abundance and Hill number over time.** The regression lines and 95% confidence intervals show the predicted relationship between time (in years) and the effects of plant species richness arthropod estimated richness (A-D), abundance (E-G), and Hill number (I-L). These were both derived from model (3), in which time was included as a numerical variable. To make the effects of plant species richness derived from model (2) comparable across years and independent of the absolute differences in arthropods between years (see Figure S3), the PSR slopes were corrected by dividing them by their respective year´s average value. The data are shown for herbivores (blue) and predators (orange) in both seasons (triangles for spring and dots for summer).
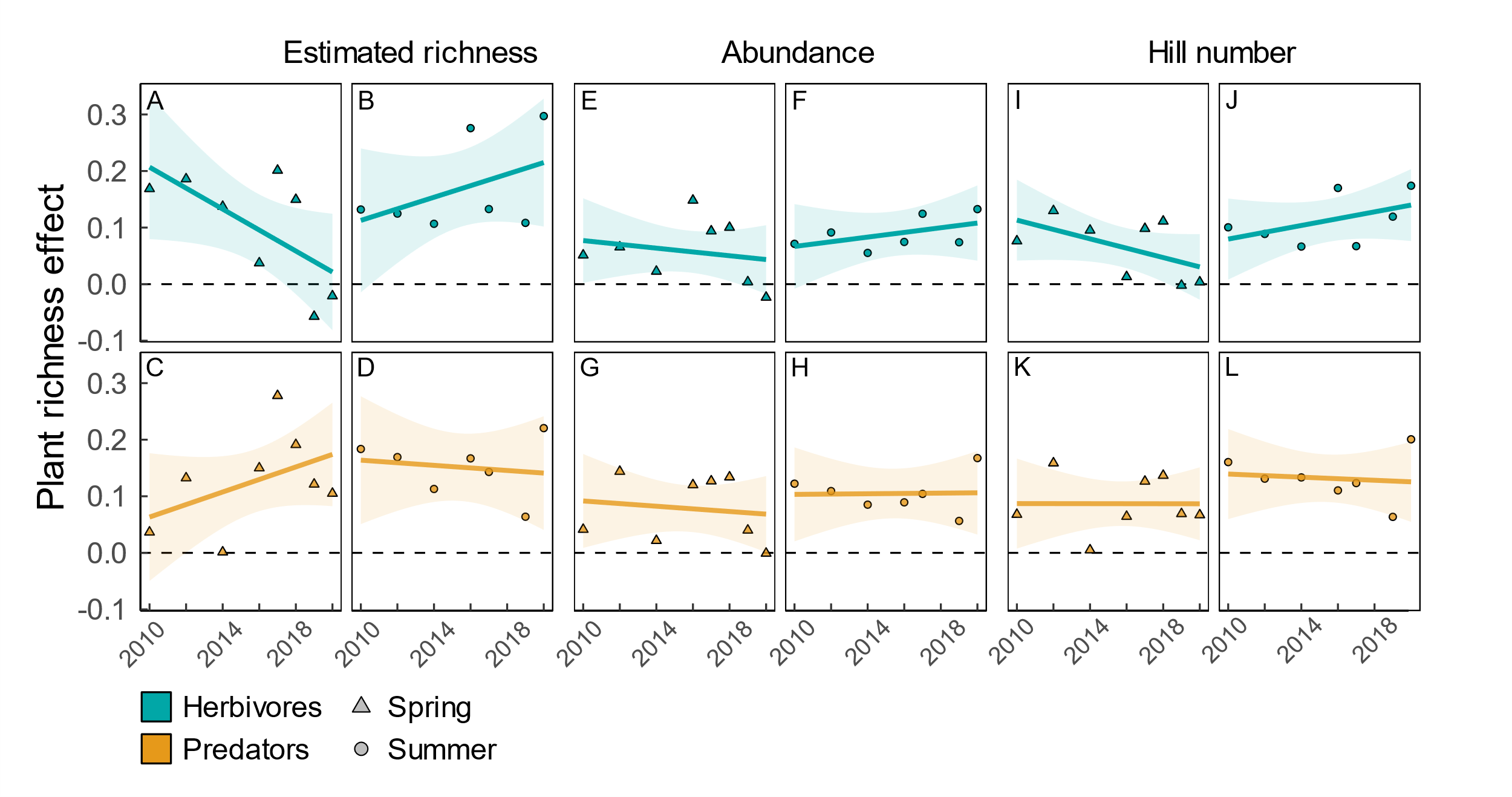

**Figure S8: Plant species richness effects on biomass ratios in multiple years.** Regression lines show the predicted relationship between plant species richness and the biomass ratio of arthropods and plants (A), herbivores and plants (B), and predators and herbivores (C) for each sampling years. Regression lines were derived from model (2), where time was included as a factor, to allow for year-to year variation in the effects of plant species richness. Arthropods and plants were sampled between 2010 and 2020. All biomass ratios and plant species richness were log-transformed prior to the analyses.

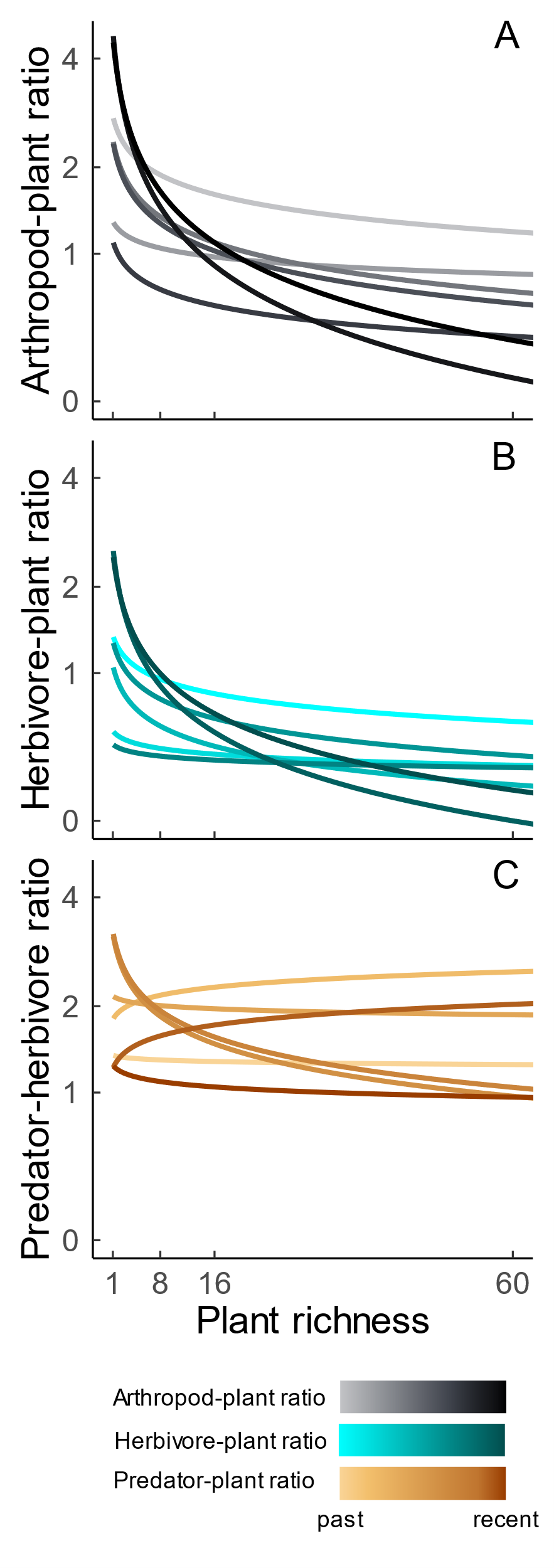

**Figure S9**: **Plant species richness effects on biomass ratios between 2010 and 2020.** Arthropod and plant biomass values are given in mg and g per m², respectively. Solid lines show main effects of plant species richness (plus confidence intervals) on biomass ratios derived from model (2). Dotted and dashed lines show plant species richness effects for spring (May) and summer (July) sampling, respectively. In 2018 no main effects could be generated, as only arthropod data from May sampling were available.

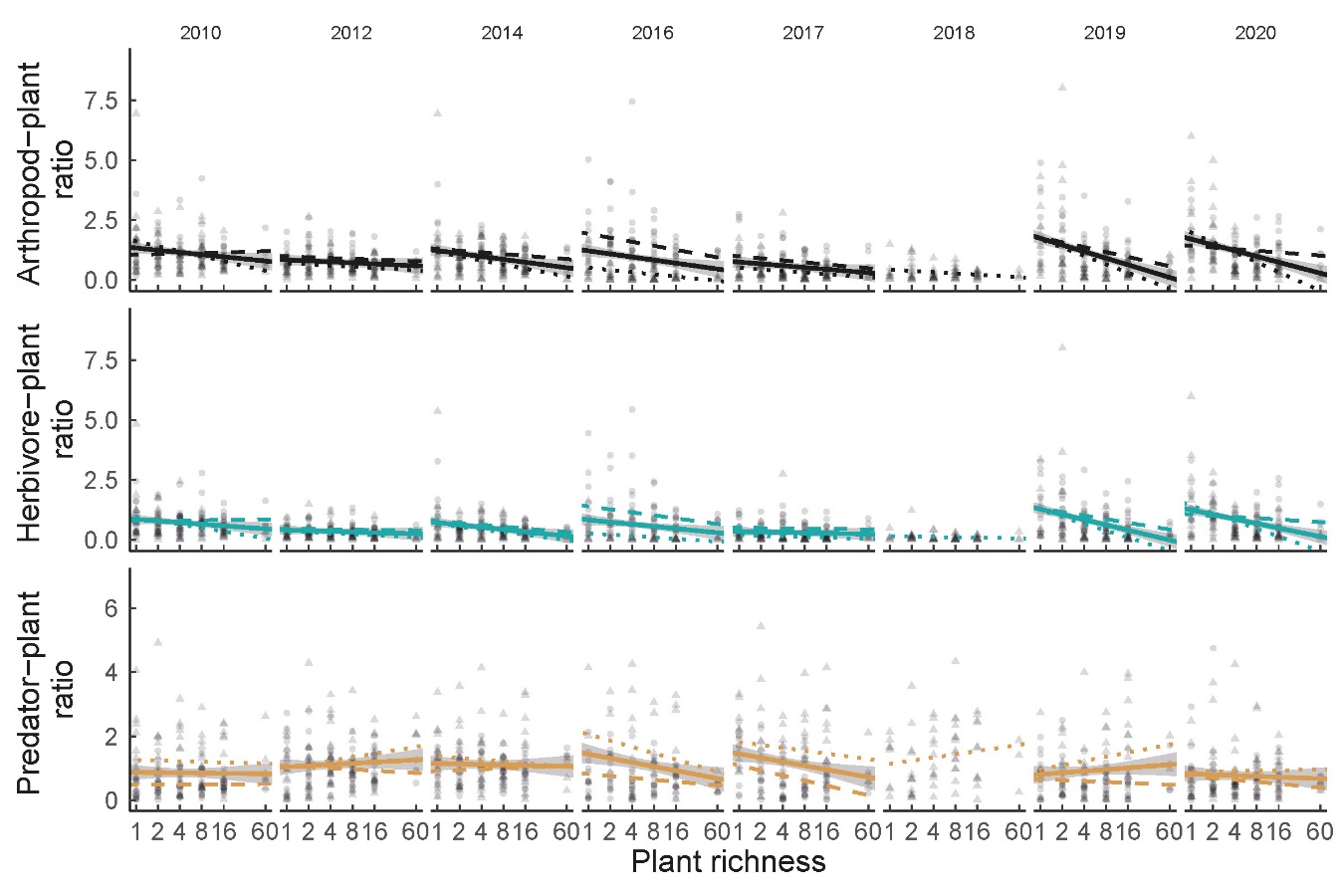

**Figure S10**: **Plant species richness effects on predation and herbivory between 2010 and 2022.** Solid lines show main effects of plant species richness (plus confidence intervals) on percentage herbivory and predation rate derived from model (2).

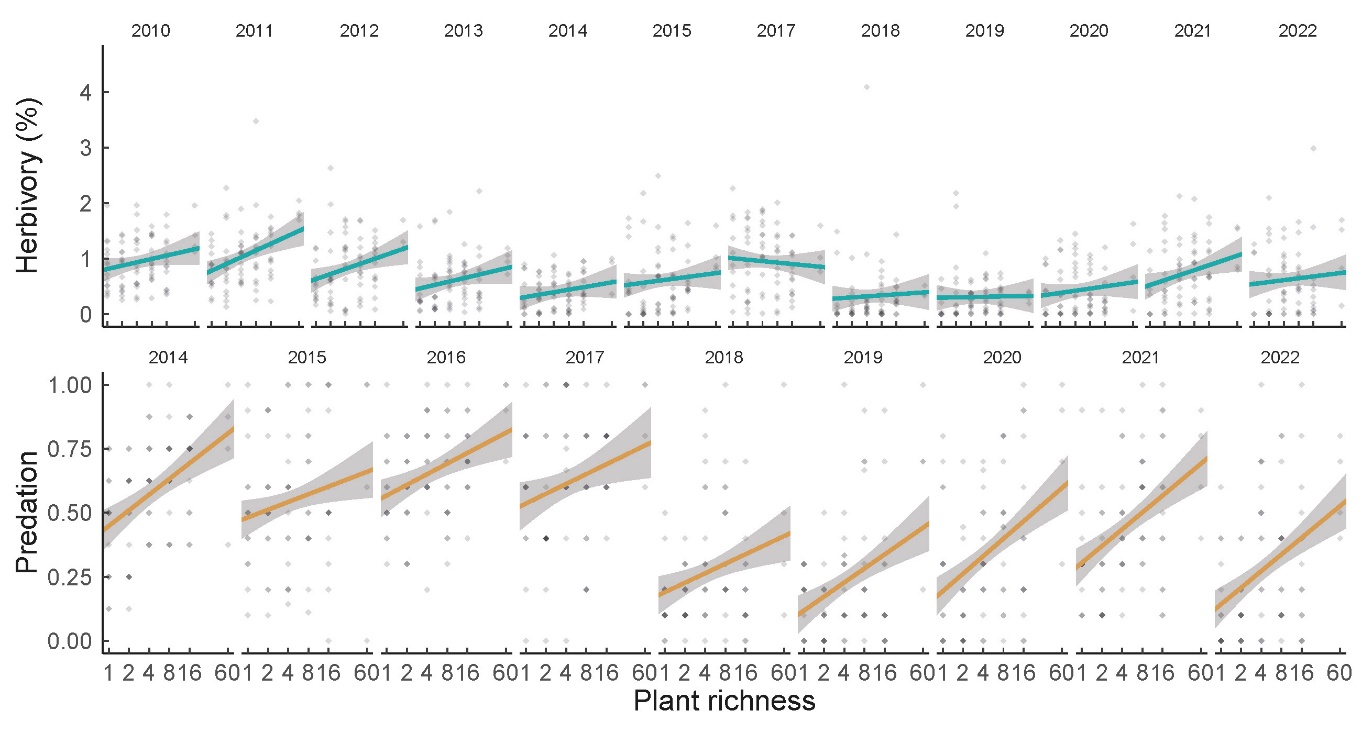

**Figure S11: Change in the effect of plant species richness on biomass ratios over time.** The regression lines and 95% confidence intervals show the predicted relationship between time (in years) and the effects of plant species richness on biomass ratios of arthropods and plants (A-B), herbivores and plants (C-D), and predators and herbivores (E-F). These both were derived from model (3), in which time was included as a numerical variable. To make the effects of plant species richness derived from model (2) comparable across years and independent of the absolute differences in arthropods between years (see Figure S8), the PSR slopes were corrected by dividing them by their respective year´s average value. Plant species richness effects were calculated separately for spring (triangles) and summer (dots).

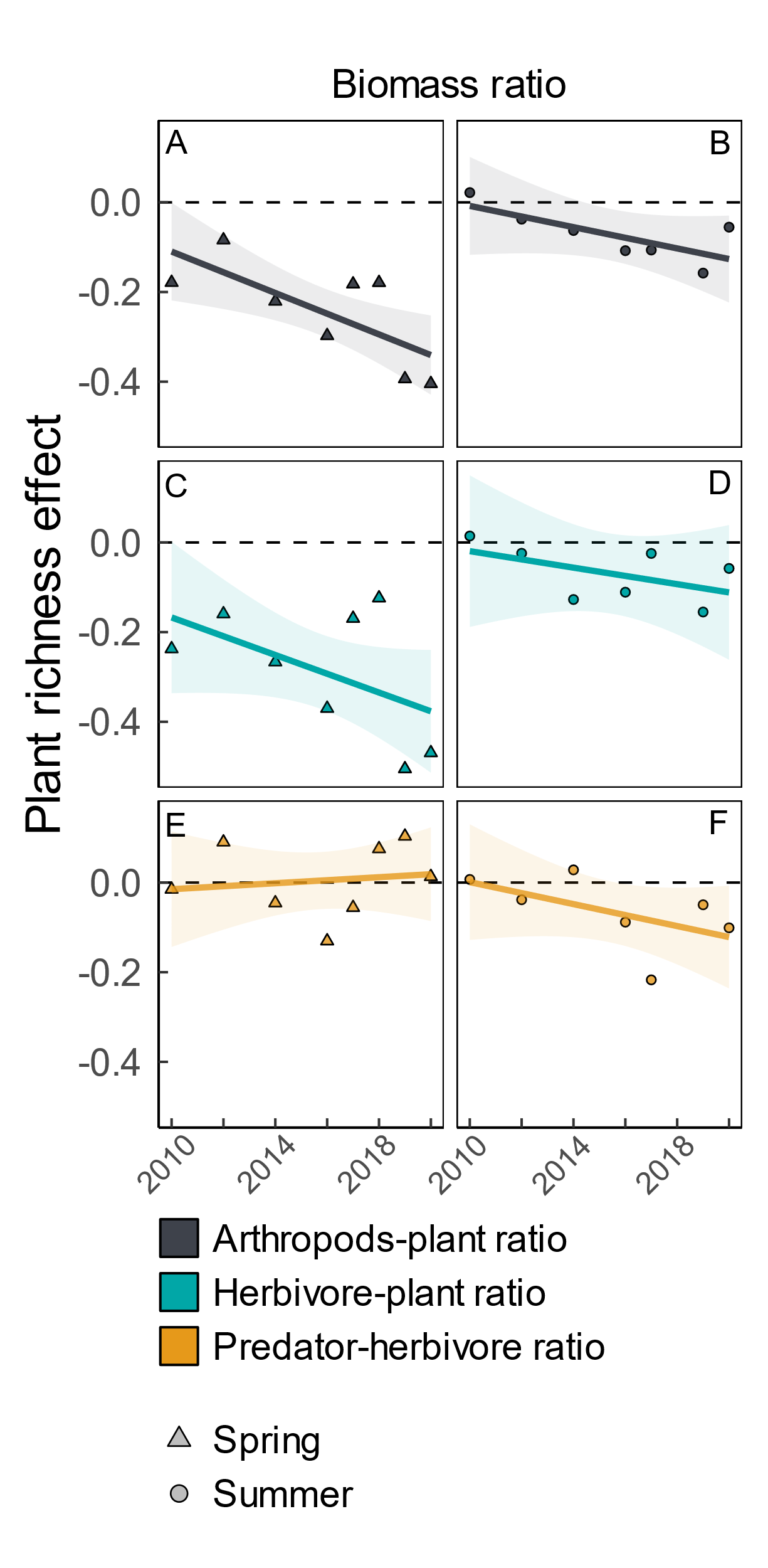

**Figure S12: Proportion of target plant species covered by the species included in the herbivory assessment.** The boxes illustrate the distribution of plant coverage across the gradient in plant species richness (top graph) and over time (bottom graph). The black lines within boxes represent median values, the boxes indicate interquartile ranges (IQR), and the whiskers show the minimum and maximum values within 1.5 x IQR. Individual raw data points are overlaid to show the data distribution. The red line highlights the median value across all plant species richness levels (top graph) or sampling years (bottom graph). Coverage was calculated by summing the cover of all species for which herbivory was assessed, then calculating their proportion of the total plant cover on the plot.

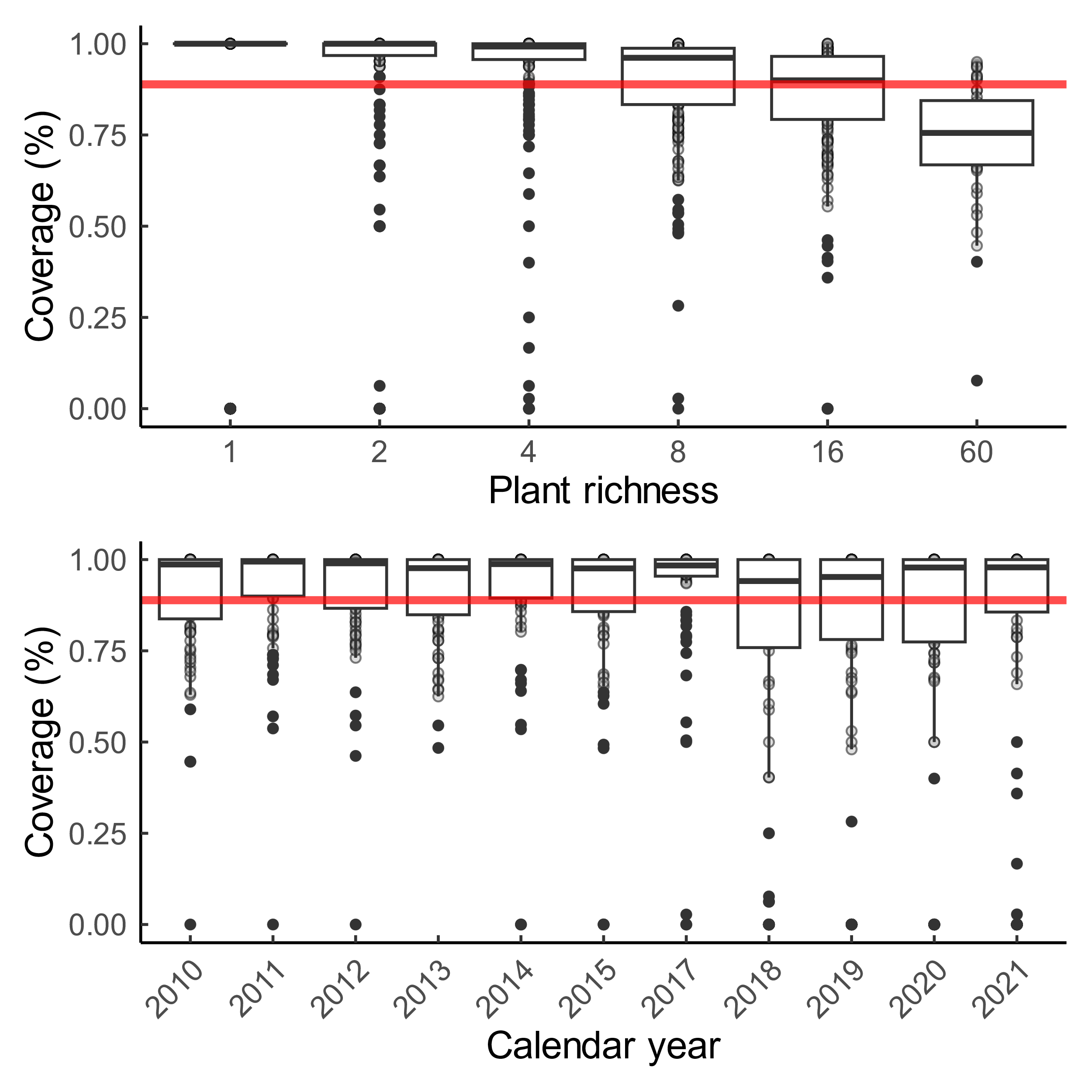
